# FoTO1 orchestrates Taxol biosynthesis through catalytic and non-catalytic mechanisms

**DOI:** 10.64898/2026.03.21.713420

**Authors:** Chloe Wick, Anish Somani, Jack Chun-Ting Liu, Sumudu S Karunadasa, Shou-Ling Xu, Polly M. Fordyce, Conor James McClune, Elizabeth S. Sattely

**Author notes:** Corresponding author: Elizabeth S. Sattely, Conor J. McClune.

## Abstract

Taxol is a blockbuster chemotherapeutic derived from the Pacific Yew tree. Recent work in our group has identified a complete pathway to baccatin III, a key intermediate, that hinges on a novel accessory protein, Facilitator of Taxane Oxidation (FoTO1). This protein dramatically improves yield and alters enzyme product profiles when reconstituting the Taxol pathway in *N. benthamiana*. FoTO1 has been shown to act early in the biosynthetic pathway improving the yields of product generated by the combination of a plastidial diterpene synthase (taxadiene synthase) and an endoplasmic reticulum (ER) localized cytochrome P450 (T5-alpha-hydroxylase). Here, we show that FoTO1 is an enzyme capable of converting taxadiene-(4),5-epoxide, the likely product of T5αH oxidation, into taxadien-5α-ol. FoTO1 is also functional in yeast, resolving a key bottleneck for development of a bioproduction route to Taxol in this host. Targeted mutagenesis of key catalytic residues in FoTO1 abrogates function *in vitro* but not *in planta*, suggesting non-catalytic contributions of FoTO1 to the taxane pathway. A combination of proximity labelling, bimolecular fluorescence complementation assays, and co-immunoprecipitation studies revealed that FoTO1 interacts with and organizes various P450s in the Taxol pathway. These approaches highlight the importance of both FoTO1’s catalytic and non-catalytic functions in improving yields in the early Taxol pathway. Beyond Taxol biosynthesis, FoTO1 boosts yields for diverse diterpene pathways from across phylogeny, suggesting a general role of this protein class in mediating metabolism across the plastid and ER in plants.

## Introduction

Diterpenes produced in plants are a key source of many medicinal compounds^1,2^. A lead example is paclitaxel (trade name Taxol), a chemotherapeutic used in the treatment of breast, ovarian, and lung cancer, that is produced by the yew tree (*Taxus spp*.)^3,4^. Taxol production is challenging due to its chemical complexity and low natural abundance in the yew tree^5–7^. Notable efforts have led to synthetic routes for production of Taxol; however these have not been scaled for manufacture to date^8–11^. Currently, Taxol manufacture is still dependent on the extraction of metabolites from yew tissue^12^. Thus, the elucidation of the biosynthetic route to this molecule has been a major focus for the development of a bioproduction process^13–15^. Recent efforts in our lab and others have culminated in the discovery of a complete set of genes that enable reconstitution of late stage intermediates in Taxol biosynthesis, including baccatin III^16 18^

From Taxol to gibberellin, diterpene biosynthesis pathways consistently follow the same general architecture. Pathways begin with diterpene synthases that usually localize to the plastid^19,20^. These enzymes catalyze the formation of the core diterpene scaffold, representing the first committed steps in most known diterpene pathways. The resulting scaffolds are subsequently modified by tailoring enzymes, including cytochromes P450 enzymes anchored in the endoplasmic reticulum (ER), as well as 2-ODDs and acetyltransferases^21^. This compartmentalization of the pathway between the chloroplast and the ER suggests a need to coordinate biosynthesis across these organelles.

In the case of the Taxol pathway, taxadiene synthase (TDS) produces the diterpene scaffold taxadiene, which is then thought to be a substrate for oxidation by the cytochrome P450 taxadiene-5α-hydroxylase (T5αH)^22^. However, in attempts to reconstitute this pathway, T5αH catalysis primarily results in the formation of multiple undesired side products *in vitro* and in heterologous hosts, especially 5(12)-oxa-3(11)-cyclotaxane (OCT, **2a**) and/or 5(11)-oxa-3(11)-cyclotaxane (iso-OCT, **2b**), which has been the source of a long-standing bottleneck in the reconstitution of Taxol biosynthesis^13,23,24^. Interestingly, these byproducts are observed only in reconstitution systems but not in the native *Taxus* tree, suggesting a missing relevant protein or mechanism that enables specificity by T5αH catalysis^17^. In previous work, we discovered that a previously uncharacterized protein, FoTO1, improves product yields by an order of magnitude and reduces side product formation, although it is not essential for pathway reconstitution in the heterologous host *N. benthamiana*^17^. FoTO1 is a member of the NTF2-like superfamily of proteins which contains over 450,000 members (InterPro). This versatile family contains proteins found across bacteria, fungi, and plants with diverse functions, including enzymatic, transport, and scaffolding roles^25,26^. In plants, NTF2-like proteins in *Arabidopsis thaliana* have been shown to be involved in the nuclear transport of GTPase Ran and NTF2 proteins in *Medicago sativa* and *Glycine max* are involved in drought tolerance^27–29^. However, to our knowledge, no NTF2-like superfamily members have previously been shown to play a role in metabolic pathways in plants.

FoTO1 appears to influence early pathway flux by modulating the product profile of pathway enzymes. Specifically, inclusion of FoTO1 in reconstitution experiments with T5αH leads to the selective formation of the desired, productive intermediate taxadien-5α-ol and elimination of side products OCT and related compounds^17^. However, the role of FoTO1 in the Taxol pathway has remained unclear. Here, we characterize the mechanisms by which FoTO1 improves both yield and selectivity in the early Taxol pathway, resolving a critical bottleneck for Taxol bioproduction. We find that FoTO1 not only acts as an enzyme, converting a previously unverified epoxide intermediate into the correct product, but also performs additional non-catalytic roles relevant to diverse diterpene pathways. We employ a multifaceted approach that integrates *in vitro* and heterologous pathway reconstitution, and interaction studies *in planta* and with purified protein to establish that FoTO1 enhances yield improvement through two distinct mechanisms: (1) catalyzing the conversion of taxadiene epoxide to taxadien-5α-ol, (2) organizing pathway enzymes through transient protein-protein interactions. Lastly, we show that *Taxus x media* FoTO1 can be used to enhance the yields of plant diterpene pathways from diverse species. This model is supported by protein mutagenesis that decouples FoTO1’s two distinct functions; specifically, a catalytically inactive mutant still provides a substantial yield improvement. Together these data shed light on the long-standing bottleneck in Taxol biosynthesis and suggest a new role for the NTF2-like protein superfamily in coordinating multistep metabolic pathways across organelles in plants.

## Results

### FoTO1 improves selectivity and yield of taxadien-5α-ol in yeast

Previously, we demonstrated that FoTO1 leads to a robust improvement in the yield of taxadien-5α-ol and elimination of side products when TDS and T5αH are expressed using *Agrobacterium*-mediated transient DNA delivery into *Nicotiana benthamiana* (tobacco) leaves^17^. This yield improvement propagates to the final products (e.g. baccatin III) of an engineered taxane pathway and resolves a major outstanding bottleneck to Taxol bioproduction. However, two key questions remain unresolved: how does FoTO1 function, and is it broadly useful for metabolic engineering of diterpene pathways? To address whether FoTO1-mediated enhancement of pathway yield requires plant specific components or simply a eukaryotic cell context, we turned to yeast as a host. Specifically, we sought to investigate whether FoTO1 is sufficient to improve yields of taxane products in this host. We transformed a FoTO1-expressing plasmid into a *Saccharomyces cerevisiae* yeast strain that harbors integrated TDS and T5αH^30^. In the absence of FoTO1, the yeast strain harboring TDS and T5αH produced taxane rearrangement products, including **2d** (a minor product in tobacco known to be prevalent in yeast), OCT, and trace amounts of iso-OCT and taxadien-5α-ol^24^. However, when gene expression of FoTO1 was induced in this strain, a significant increase in the yield and selectivity for taxadien-5α-ol was observed (**Figure 1a**). These data show that the effects of FoTO1 do not require a plant-cell specific environment to increase taxadien-5α-ol production and selectivity. Furthermore, these results pave the way for Taxol biomanufacture in industrial yeast strains.

**Figure 1:**
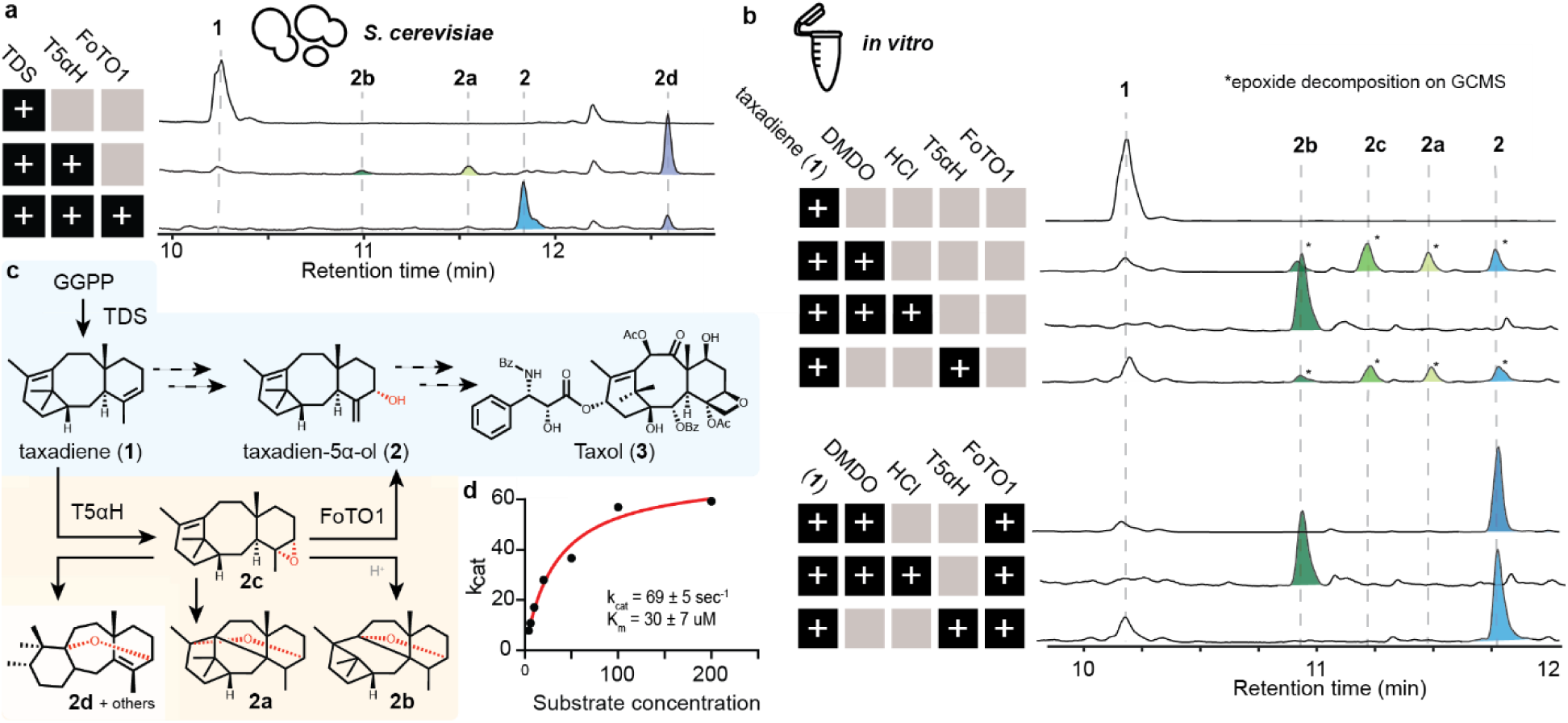
Biochemical characterization of FoTO1 and identification of early epoxide intermediate as a substrate. (a) GCMS total ion chromatogram (TIC) of *S. cerevisiae* strains integrated with the early Taxol biosynthetic pathway enzymes TDS; TDS and T5αH; or TDS and T5αH transformed with FoTO1-carrying plasmids. TICs representative of three independent experiments. (b) GCMS TIC of *in vitro* reactions pairing taxadiene with enzymatic (T5αH from yeast microsomes) and chemical (Dimethyldioxirane (DMDO)) oxidants, with or without purified FoTO1 added after conversion of oxidative reaction. Enzymatic activity of purified FoTO1 was also tested after DMDO oxidation followed by acid isomerization into degradation products. The scale of the y-axis is consistent across reactions, representing the products. TICs representative of three independent experiments. (c) Proposed chemical scheme of the catalysis from taxadiene to taxadien-5α-ol by reactions with T5αH and FoTO1. (d) kcat and Km values of FoTO1 for the reaction of the taxadiene-4(5)-epoxide substrate, calculated using 7 different concentrations of epoxide substrate (4μM - 200μM) in liposomes. Catalytic activity was estimated through GCMS quantification of substrates and products at multiple timepoints, with extractions normalized to internal standard. The graph shown here is representative of three independent experiments.

### FoTO1 catalyzes the conversion of an epoxide intermediate to the desired product taxadien-5α-ol

Considerable effort has been devoted to understanding the production of OCT and iso-OCT – unintended side products generated during the initial T5αH-catalyzed oxidation of the taxadiene scaffold – across multiple reconstitution systems. Previously, we structurally characterized several of these side products and found that many represent rearranged isomers and overoxidation products^31^. Several different mechanisms for this apparent catalytic promiscuity of T5αH have been proposed. Initial papers suggested that the formation of multiple products was inherent to the enzyme, though other proposals instead suggest that a taxadiene-4(5)-epoxide intermediate is formed first, and then degrades to the side products in solution^24,32^. Since inclusion of FoTO1 greatly reduces off-pathway products in yeast and plants, we hypothesized that this protein might be able to enzymatically convert a potential epoxide intermediate into taxadien-5α-ol. Although FoTO1 homologs are not known to contribute to metabolism in plants, enzymes with a similar fold have been characterized in polyketide biosynthesis and epoxide catabolism pathways from bacteria^33,34^. To test whether FoTO1 has a catalytic role in resolving an unstable taxane pathway intermediate, we chemically synthesized taxadiene-4(5)-epoxide using dimethyldioxirane (DMDO), as previously reported^35^. We observed no instability when this molecule is stored in pyridine-d5 at room temperature over 3 days prior to analysis using NMR (**Supplementary Fig 1-2**). However, our analysis by gas chromatography mass spectrometry (GC-MS) of the same taxadiene-4(5)-epoxide sample immediately following synthesis revealed multiple peaks that appear to correspond to taxadien-5α-ol (**2**), the undesired intermediates OCT (**2a**) and iso-OCT (**2b**), as well as a fourth peak which we proposed to be taxadiene-4(5)-epoxide (**2c**) (**Figure 1b**).

Given the purity of the epoxide as indicated by NMR, we surmised that the multiple peaks observed upon GC-MS analysis likely result from thermal degradation of the epoxide during injection. Strikingly, addition of purified FoTO1 (obtained through heterologous expression and purification from *E. coli*) to the synthetic taxadiene-4(5)-epoxide *in vitro* yielded a single taxadien-5α-ol peak with no observable degradation products present upon GC-MS analysis (**Figure 1b, line 5**). These results suggest a catalytic role for FoTO1, with taxadiene-4(5)-epoxide serving as its direct, enzymatic substrate. Kinetic analysis of purified FoTO1 protein incubated with taxadiene-4(5)-epoxide in reconstituted liposomes revealed that FoTO1 turns over its epoxide substrate with a kcat of 69 ± 5 sec^−1^ and a Km of 30 ± 7 µM (**Figure 1c; Supplementary Fig 3**). To investigate whether FoTO1 is specific toward taxadiene-4(5)-epoxide or is also able to isomerize other side-products like OCT and iso-OCT, we converted taxadiene-4(5)-epoxide to iso-OCT using 1M HCl following a previously reported method and subjected it to FoTO1 *in vitro*^35^. We found that incubation with FoTO1 does not result in the consumption of iso-OCT or formation of new isomerized products, suggesting that FoTO1 activity is specific to taxadiene-4(5)-epoxide (**Figure 1b, line 6**). Together, these data provide evidence that OCT-related metabolites are indeed derailed pathway intermediates unlikely to be relevant to taxane biosynthesis in the Pacific Yew. Furthermore, FoTO1 acts as a catalyst to enable productive resolution of pathway intermediates towards elaborated taxanes.

We next sought to determine a possible biosynthetic origin for the taxadiene-4(5)-epoxide. When we incubated taxadiene with T5αH-expressing microsomes from yeast and compared the GC-MS spectra to those of the NMR-verified epoxide in organic solvent, we observed similar results. Both spectra had matching ratios of OCT, iso-OCT, taxadien-5α-ol and a small but detectable hallmark epoxide peak (**Figure 1b, line 4 vs line 2**). These findings align with previous reports showing that the GC-MS spectra of taxadiene oxidized by T5αH and taxadiene oxidized by DMDO are remarkably similar^24^. Consequently, we found that adding FoTO1 to taxadiene incubations containing T5αH-expressing microsomes also substantially boosted taxadien-5α-ol production (**Figure 1b; Supplementary Fig 4-5)**.These data supports a model where the taxadiene-4(5)-epoxide is the direct product of T5αH, with the apparent promiscuity arising from the formation of multiple degradation products during analysis (either by incubation in acidic extracts or thermal injection on GC-MS). To see whether we could observe this epoxide product *in planta*, we infiltrated a low titer of *A. tumerificans* (OD of 0.04) encoding T5αH alongside *Agrobacterium* strains encoding taxadiene synthase and genes boosting diterpene metabolic flux *in N. benthamiana* leaves. On close analysis, we are able to identify the hallmark epoxide peak concurrently with high amounts of rearrangement products (**Supplementary Fig 6-7**). The high levels of iso-OCT and OCT *in planta* suggest that the epoxide undergoes some decomposition *in planta* prior to degradation on the GCMS. Thus, these *in vitro* and *in planta* data together indicate that the direct product of T5αH oxidation of taxadiene is taxadiene-4(5)-epoxide, and suggest FoTO1 then captures and resolves this intermediate to taxadien-5α-ol prior to degradation.

### Residues D149, D68, and Y60 are critical for FoTO1 enzymatic activity

To further characterize the catalytic activity of FoTO1, we sought to identify active-site residues and generate catalytically inactive FoTO1 mutants, through comparison with structurally homologous enzymes. Several bacterial and fungal enzymes share a similar NTF2 fold with FoTO1, including two epoxide hydrolases, two polyether synthases, and the TerC dehydratase which all have epoxide substrates or intermediates^33,34,36–38^. Although the primary sequences of these proteins vary considerably (19% amino acid identity between LEH and FoTO1), their folds are conserved and similar to the AlphaFold-predicted structure of FoTO1.

Pairwise comparison between the selected NTF2 enzymes and FoTO1 provided insight into FoTO1’s catalytic mechanisms. The two epoxide hydrolases and TerC act on cis-epoxides fused to 6-membered rings, substrates with similar geometries to taxadiene-4(5)-epoxide. To facilitate cis-epoxide ring-opening, the active sites of these three enzymes all contain a spatially conserved aspartate annotated to function as a general acid catalyst. Structural alignment of these three structures with the predicted structure of FoTO1 revealed that these catalytic aspartates overlap with D149 in FoTO1 (**Figure 2a; Supplementary Fig 8**). The NTF2-like polyether synthases Lsd19 and MonBI act on epoxides with different geometries to FoTO1, however they both conduct similar base-catalyzed intramolecular rearrangement to open the epoxide. A key conserved aspartate functions as a catalytic base, which overlaps with D68 in the predicted FoTO1 structure (**Figure 2a, Supplementary Figure 8**).

**Figure 2:**
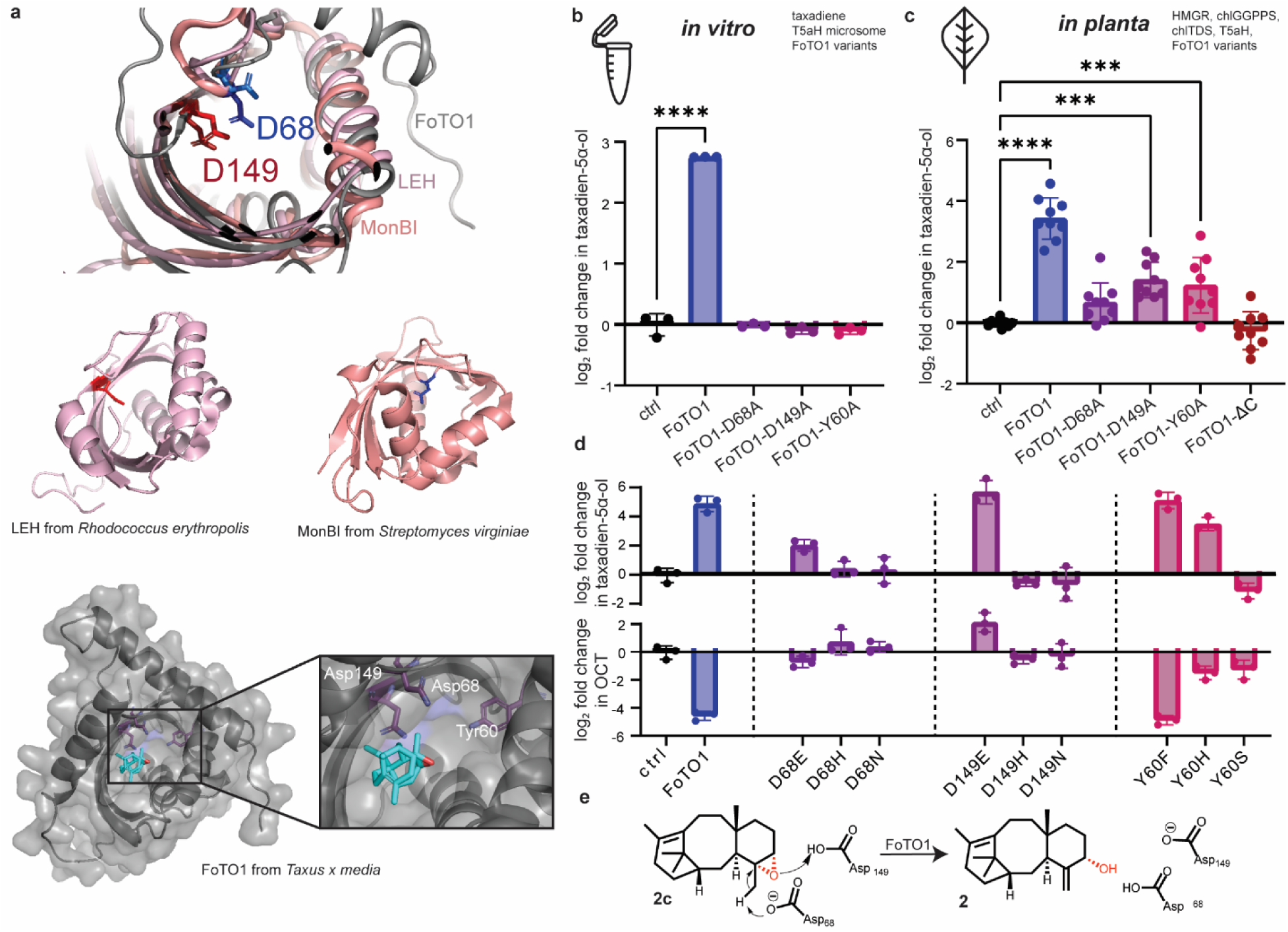
Point mutants of FoTO1 that abrogate activity *in vitro* still retain some activity when tested *in planta*. (a) Proposed active site of FoTO1 (generated by AlphaFold3) aligned with the limonene-1,2-epoxide hydrolase (LEH; *R. erythropolis*) and MonBI (*S. virginiae*) crystal structures. Proposed catalytic residues in FoTO1 and their counterparts in LEH and MonBI are highlighted. Structures of LEH and MonBI with highlighted residues D132 and D37, respectively, are displayed below the alignment of all three structures and the AlphaFold predicted structure of FoTO1 with the proposed epoxide substrate docked using DiffDock is shown in grey at the bottom. The three residues hypothesized to be essential for catalysis are highlighted in purple. (b) Quantification of the log2 fold change of taxadien-5α-ol when point mutants of FoTO1 purified from *E. coli* are incubated with taxadiene *in vitro*. Values ± SD of three independent experiments (∗p ≤ 0.05, ∗∗p ≤ 0.01, ∗∗∗p ≤ 0.001, ∗∗∗∗p ≤ 0.0001 by ordinary one-way ANOVA and not shown if insignificant (p > 0.05)). (c) Quantification of the log2 fold change of taxadien-5α-ol when point mutants of FoTO1 are transiently expressed in *N. benthamiana* leaves along with boost and early pathway enzymes (HMGR, GGPPS, chlTDS, T5αH). mCherry (ctrl) is used instead of FoTO1 as a negative control and wild-type FoTO1 is used as a positive control. Fold change is calculated as the ratio of the measured peak area of taxadien-5α-ol in a sample divided by the average measured peak area of taxadien-5α-ol in the absence of FoTO1 (control) of that run. Values ± SD of nine independent experiments (∗p ≤ 0.05, ∗∗p ≤ 0.01, ∗∗∗p ≤ 0.001, ∗∗∗∗p ≤ 0.0001 by ordinary one-way ANOVA and not shown if insignificant (p > 0.05)). (d) Quantification of OCT and taxadien-5α-ol when point mutants of FoTO1 are transiently expressed in *N. benthamiana* leaves along with boost and early pathway enzymes (HMGR, GGPPS, cytTDS, T5αH). Values ± SD of three independent experiments (∗p ≤ 0.05, ∗∗p ≤ 0.01, ∗∗∗p ≤ 0.001, ∗∗∗∗p ≤ 0.0001 by ordinary one-way ANOVA and not shown if insignificant (p > 0.05)). (e) Proposed enzymatic mechanism for FoTO1 catalytic activity involving two catalytic aspartate residues, Asp68 and Asp149, which are indispensable for catalysis.

Computational docking also independently suggested key catalytic roles for the two aspartates. Docking of either the epoxide substrate or the taxadien-5α-ol product into the predicted FoTO1 structure placed the ligands in the binding pocket proximal to the two aspartates (**Figure 2a, Supplementary Fig 9**). Upon docking of taxadien-5α-ol, the two aspartates were observed to be within **3 Å** of the protons transferred during epoxide opening and allylic alcohol formation, with D149 appearing to function as a catalytic acid and D68 as a catalytic base, agreeing with predictions from homologous enzymes (**Supplementary Fig 9**).

We tested the importance of these amino acids in FoTO1 catalysis through site-directed mutagenesis and found that mutation of either D149 or D68 to alanine completely abrogated the ability of FoTO1 to transform taxadiene-4(5)-epoxide into taxadien-5α-ol *in vitro* (**Figure 2b**). To identify other residues important in catalysis, an additional 14 residues in the predicted inner pocket of FoTO1 were screened. Only mutation of Y60 had a significant effect on the amount of taxadien-5α-ol produced (**Figure 2-b, Supplementary Fig 10-11**). These data suggest that the residues D149, D68, and Y60 in FoTO1 are directly involved in promoting isomerization (or binding) of the taxadiene-4(5)-epoxide to generate the productive pathway intermediate taxadien-5α-ol.

To further probe the role of acidic residues in the active site, we also examined a set of FoTO1 mutants with conservative substitutions, replacing aspartate with asparagine or glutamate. When tested in *N. benthamiana,* plants expressing FoTO1 mutants D68N and D149N had significant accumulation of iso-OCT and OCT rearrangement products, similar to what was observed with D68A and D149A. However, mutation to D68E and D149E resulted in the partial recovery of catalytic activity and a reduction in the observed rearrangement products (**Figure 2d**). We also generated a Y60F mutant with retained aromaticity. Y60A abolished catalytic function but Y60F did not, suggesting a potential structural role for this residue (**Figure 2b,d, Supplementary Fig 11**). Taken together, these point mutants of FoTO1 allowed for the specific analysis of the catalytic functions of FoTO1 while still maintaining the general structural features of the protein.

### FoTO1 has additional non-catalytic properties in planta

Previously, we showed that T5αH and FoTO1 directly interact using co-immunoprecipitation and *in vitro* microscale thermophoresis binding experiments, suggesting that FoTO1 may have additional functions in a plant cell beyond catalysis^17^. To examine the contributions of these possible non-catalytic roles to the enhanced yields of taxane pathway intermediates (e.g. through scaffolding or protein-protein interactions), we leveraged our catalytically inactive FoTO1 mutants (D149A, D68A, Y60A) for experiments *in planta.* These mutants were tested in the context of other pathway enzymes, cellular components, and intact organelles for a role in pathway flux. As described above, these catalytically inactive mutants showed no yield enhancement when tested with our *in vitro* assay system (**Figure 2b**). To test the effect of FoTO1 on the early Taxol pathway *in planta*, we used *Agrobacterium*-mediated transient DNA delivery to express, in *Nicotiana benthamiana*, boost enzymes (HMGR, GGPPS) that increase the pool of taxadiene precursors alongside the Taxol pathway enzymes taxadiene synthase (TDS) and T5αH^39–41^. Unlike the complete lack of function observed in our *in vitro* assays, expression of the FoTO1 catalytic mutants D149 or Y60A in combination with T5αH, TDS, and boost enzymes *in planta* resulted in a reproducible 3-fold increase in taxadien-5α-ol and a slight, but non-significant increase in relevant side products (iso-OCT, OCT) – relative to control experiments without FoTO1 (**Figure 2c, Supplementary Fig 10**). These data suggest that FoTO1 could have additional roles *in planta* beyond catalysis, observable even in a heterologous plant host.

### FoTO1 interacts with multiple Taxus P450s

Given the significant increase in taxadien-5α-ol production by FoTO1 mutants *in planta* but not *in vitro*, we wanted to further investigate potential non-catalytic contributions of FoTO1. Non-catalytic contributions of biosynthetic enzymes have recently been emphasized in the reconstitution of a select few plant specialized metabolic pathways. Lead examples include the formation of metabolons in cyanogenic glucoside dhurrin biosynthesis and steroidal glycoalkaloid biosynthesis^42–46^. Additional examples have also emerged in phenylpropanoid metabolism in plants and glycosylation in mammalian systems. Central to these examples is the role protein-protein interactions play in guiding metabolic flux through enzymatic steps. Here, we were curious about FoTO1’s ability to interact with other enzymes besides T5αH in the Taxol biosynthetic pathway to orchestrate a potential metabolon that could channel products and improve yields. First, we leveraged the proximity labeling system TurboID which allowed us to simultaneously investigate infiltrated *Taxus* enzymes and native *N. benthamiana* proteins^47,48^. TurboID is an engineered, promiscuous biotin ligase that biotinylates proteins within a ∼10 nm radius (**Figure 3a**)^48^. To investigate which proteins localize near FoTO1, select Taxol-associated enzymes, including enzymes that act directly upstream and downstream of FoTO1 (T5αH, TAT, T10βH, DBAT, T13αH) as well as later pathway enzymes (T2αH, PCL, PAM) were co-expressed in *N. benthamiana* along with either a TurboID-FoTO1 fusion or a TurboID-only control (**Figure 3b, Supplementary Table 1**). Plastidial TDS and an enzyme to boost the production of taxadiene precursors (chlGGPPS) were also included as well as cytochromes P450 from unrelated terpene pathways in three other species. Tandem mass spectrometry analysis revealed significant enrichment of all Taxol pathway P450 enzymes—including T5αH, T10βH, T13αH, and T2αH—in the TurboID-FoTO1 samples as compared to the TurboID control (**Figure 3b-c**). These data suggest FoTO1 is proximal to these proteins *in planta*, albeit in the context of a heterologous expression host. Two additional non-*Taxus* P450s from *Citrus* and *Coleus* were also enriched to similar levels as T5αH, T13αH, and T2αH (**Figure 3b-c**). In contrast, Taxol pathway enzymes not associated with the ER (e.g., TDS1, TDS2, DBAT, T9dA, DBTNBT) and control proteins such as GFP were not significantly enriched in our dataset (**Figure 3b-c**). While this approach does not provide direct evidence for FoTO1-protein complexes, it reveals that FoTO1 is generally in close proximity to many transmembrane ER proteins, including cytochromes P450 from *Taxus* and the endogenous *N. benthamiana* CPR (cytochrome-P450 reductase) (**Figure 3c-d, Supplementary Fig 12**). It is notable that FoTO1 does not appear to contain an obvious ER targeting sequence, despite the enrichment of ER-localized proteins in this dataset, with thirty-six out of the top hundred enriched proteins localizing to the ER (**Figure 3d**). In contrast, only four of the top hundred proteins labelled by the TurboID control were ER-localized (**Supplementary Fig 12**). Further analysis also revealed that sixteen of the top one hundred significantly enriched TurboID-FoTO1 hits were localized to the surface of the plastid and not found in the Turbo-only control, suggesting that FoTO1, while proximate to the ER, might also be co-localized with this organelle (**Fig 3d; Supplementary Fig 12**). Thus, our untargeted proximity labelling approach suggests a possible role for FoTO1 in coordinating interactions between organelles and enzymes in the Taxol biosynthetic pathway.

**Figure 3:**
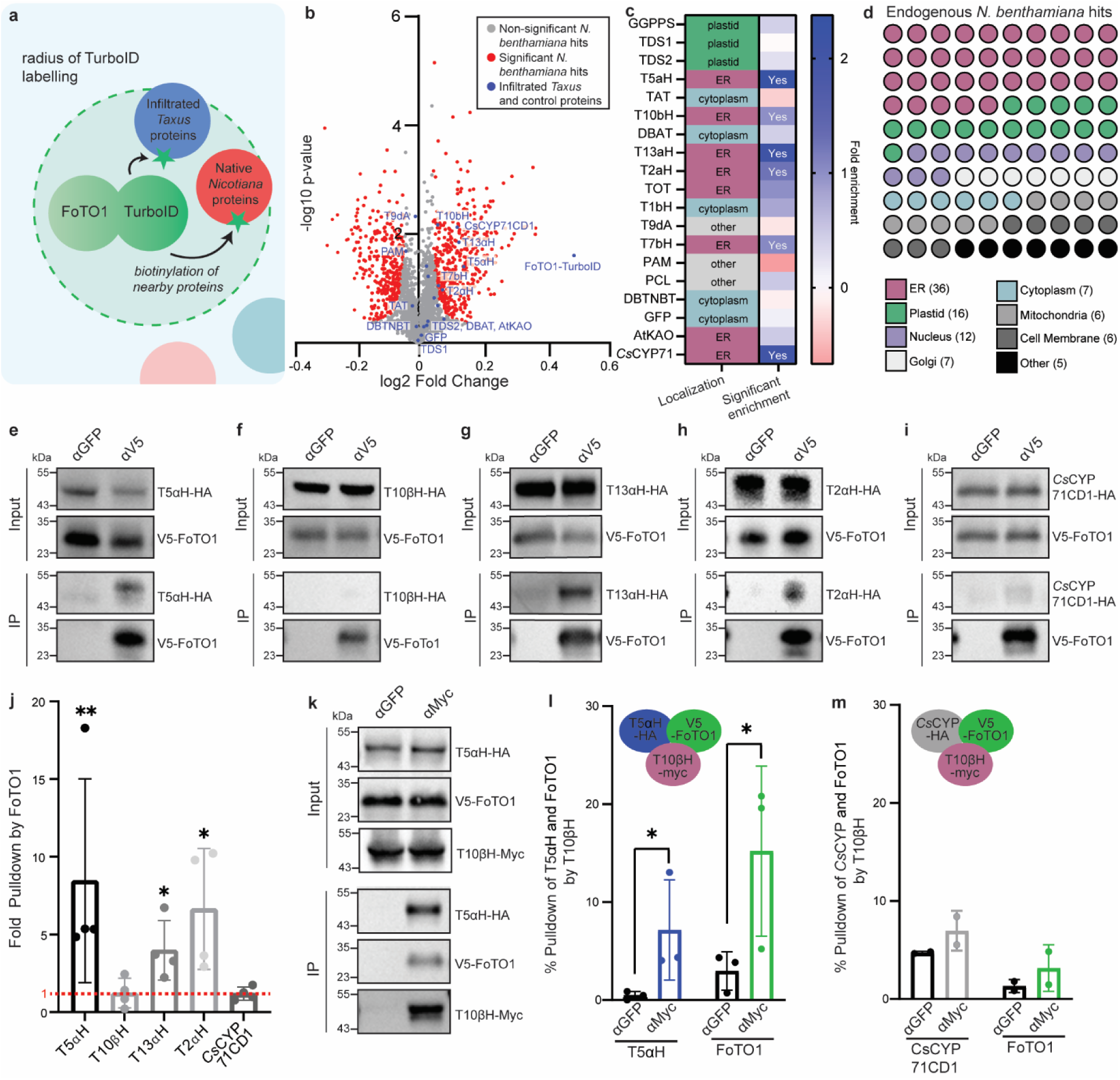
TurboID and immunoprecipitation reveals FoTO1 interacts with multiple P450s. (a) Proteins near a TurboID-FoTO1 chimera are labelled with biotin by the TurboID. These biotinylated proximal proteins are then enriched using streptavidin coated beads and identified via tandem MS/MS. (b) A volcano-scatter plot of all proteins enriched after labelling with TurboID-FoTO1. Non-significant *N. benthamiana* proteins are shown in grey while transiently expressed *Taxus* and control proteins are highlighted in blue and significantly enriched and non-enriched *N. benthamiana* proteins are shown in red. (c) Table of the infiltrated *Taxus* and control proteins displaying the localization of each protein and the fold enrichment of each protein in the TurboID-FoTO1 samples vs. TurboID control samples. Samples that are significantly enriched (p-value <0.05) are denoted in the second column. (d) Dot plot displaying the localization of the top 100 most enriched endogenous *N. benthamiana* proteins in the TurboID-FoTO1 screen. The majority of proteins localized to the ER (36), with an additional portion of the hits residing in the plastid (16) and nucleus (12). (**e-i**) Representative immunoblot analysis of the co-immunoprecipitation (co-IP) of (**d**) T5αH-HA,(e) T10βH-HA, (**f**) T13αH-HA, (**g**) T2αH-HA, (**h**) citrus CYP71CD1-HA by pulldown with V5-FoTO1 in co-infiltrated *N. benthamiana* leaves. αGFP lanes indicate internal negative control pulldowns where a GFP antibody was used for pulldown instead of a V5 antibody. Representative images of four independent experiments. Extended immunoblot of (**f**) is shown in Supplementary Fig 13a. (j) Quantification of the immunoblot analysis of the co-IP of T5αH-HA, T10βH-HA, T13αH-HA, and T2αH-HA by V5-FoTO. V5-FoTO1 was co-infiltrated into *N. benthamiana* with the indicated proteins. Fold pulldown indicates the fold difference in co-IP by the V5 antibody as compared to a control GFP antibody. Values ± SD of four independent experiments (∗p ≤ 0.05, ∗∗p ≤ 0.01, ∗∗∗p ≤ 0.001, ∗∗∗∗p ≤ 0.0001 by one-sample t test and not shown if insignificant (p > 0.05)). (k) Representative immunoblot analysis of the co-IP of T5αH-HA and V5-FoTo1 by pulldown with T10βH-Myc in co-infiltrated *N. benthamiana* leaves. αGFP lanes indicate internal negative control pulldowns where a GFP antibody was used for pulldown instead of a Myc antibody. Representative images of three independent experiments. (**l-m**) Quantification of the percent pulldown via co-immunoprecipitation (co-IP) of (**l**) T5αH-HA and V5-FoTO1 or (**m**) CsCYP71CD1 and V5-FoTO1 by T10βH-Myc. The indicated proteins were co-infiltrated into *N. benthamiana*. Values ± SD of three and two independent experiments, respectively (∗p ≤ 0.05 by Student’s t test and not shown if insignificant).

### FoTO1 participates in an enzyme complex with early P450 enzymes

While TurboID is useful for capturing transient and co-localization interactions, co-immunoprecipitation can provide direct evidence for sustained binding events. To further investigate the specificity of the interactions between FoTO1 and *Taxus* cytochromes P450, we performed a series of co-immunoprecipitation experiments using *Agrobacterium*-mediated transient DNA delivery of tagged FoTO1 and interaction partners together into *N. benthamiana* leaves. Consistent with the TurboID data, co-immunoprecipitation reproduced interactions between: FoTO1 and T5αH, FoTO1 and T13aH, and FoTO1 and T2αH. No pulldown of *Taxus* T10βH or the *Citrus* cytochromes P450 *Cs*CYP71CD1 was observed, suggesting that although these proteins are in proximity *in planta*, they may not engage in a stable interaction (**Figure 3e-j, Supplementary Fig 13**). Interestingly, addition of taxadiene during co-immunoprecipitation results in a T10βH-FoTO1 interaction, which is consistent with our previous findings in which taxadiene enhanced the co-immunoprecipitation of T5αH by FoTO1^17^. However, addition of taxadiene does not allow FoTO1 to pull down *Cs*CYP71CD1 (**Supplementary Fig 13**). Given the interaction patterns between FoTO1 and these Taxol cytochromes P450, we next asked whether Taxus cytochromes P450 directly associate with one another to establish organized ER-localized complexes

To probe the ability of P450s to organize themselves, we conducted co-immunoprecipitation assays where we analyzed the ability of T10βH (the next proposed downstream pathway cytochrome P450) to pull down both T5αH and FoTO1 when all three proteins were co-expressed together in an *N. benthamiana* leaf. Pulldown of T10βH led to the statistically significant co-pulldown of T5αH (7%) and FoTO1 (15%) (**Figure 3k-l**), indicating a relatively weak but measurable interaction. Interestingly, T10βH-T5αH interaction occurs even in the absence of FoTO1 (**Supplementary Fig 13**). However, when T5αH was substituted for a cytochrome P450 from *Citrus,* T10βH was no longer able to pull down FoTO1 nor the *Citrus* cytochrome P450, suggesting that T5αH specifically stabilizes or mediates the association of T10βH and FoTO1 *in planta* (**Figure 3m, Supplementary Fig 11**). The observed interactions between FoTO1 and multiple *Taxus* cytochromes P450s suggests the possibility that Taxol enzymes, including FoTO1, participate in transient associations, ultimately implying a multi-protein complex for taxane biosynthesis.

### FoTO1 is proximate to the ER and requires plastidial TDS for its non-catalytic functions

Given the proposed interactions between FoTO1 and various Taxol cytochromes P450, we sought to analyze FoTO1 subcellular localization and potential proximity to the cytoplasmic face of the ER *in planta*. We conducted bi-molecular fluorescence complementation (BiFC) assays where we fused FoTO1 to the N-terminal half of YFP (FoTO1-YFP_n_) and resident ER proteins to the C-terminal half of YFP (T5αH-YFP_c_) and both constructs were co-expressed in *N. benthamiana* leaves. This revealed FoTO1-YFP_n_ to be in close proximity to the cytoplasmic facing C-termini of CPR-YFP_c_ and calnexin-YFP_c_, transmembrane ER proteins^49^. Co-expression of these proteins yielded a significant increase in YFP signal compared to control experiments with YFP_c_ which exhibited no fluorescence. When we co-expressed FoTO1-YFP_n_ and YFP_c_-FoTO1, we also observed significant YFP signal, which is in line with the other epoxide-resolving NTF2s that are known to function as homodimers (**Figure 4a**)^50^. Together with the TurboID localization data, these findings suggest that FoTO1 frequently resides near cytoplasmic domains of ER membrane-bound proteins.

**Figure 4:**
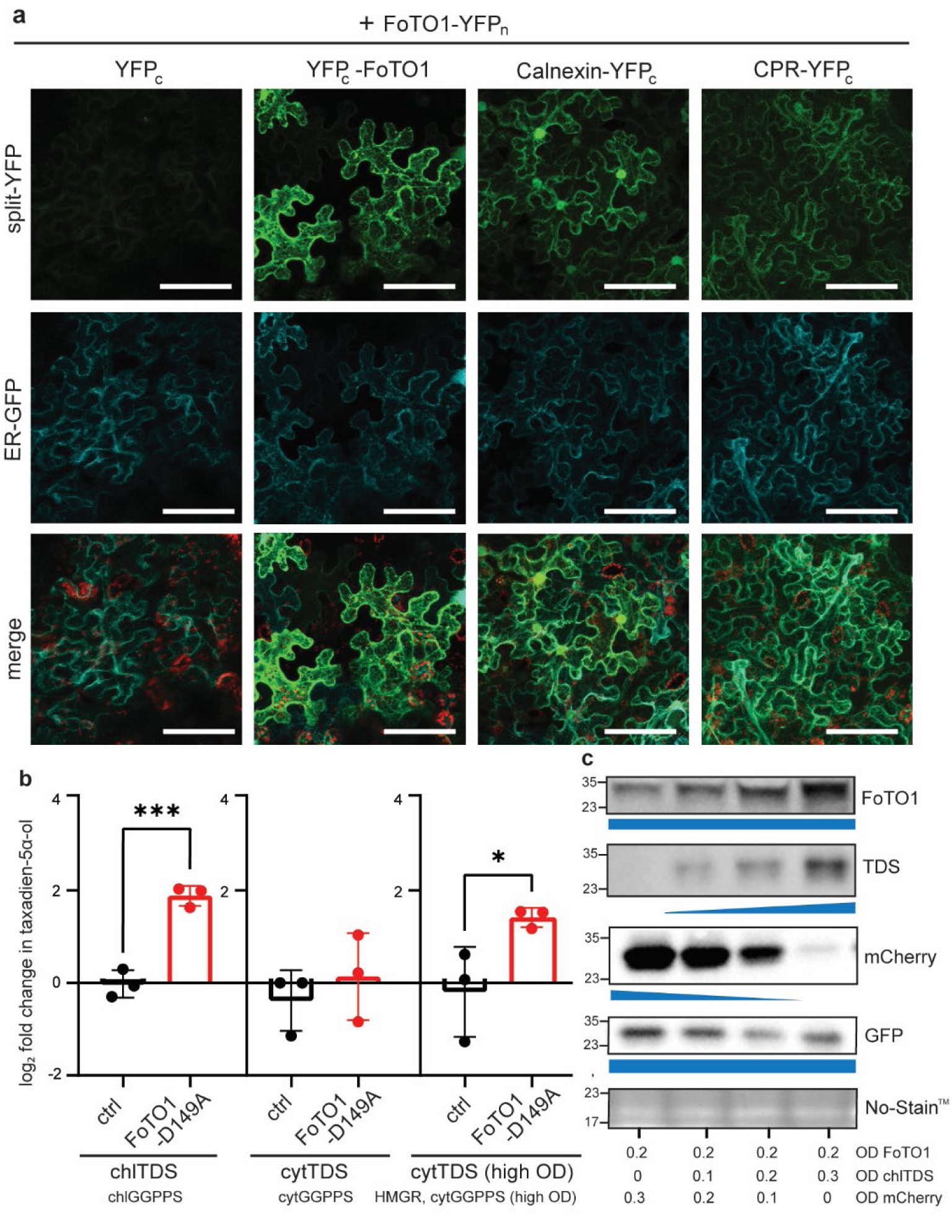
Bimolecular fluorescence complementation to analyze localization of FoTO1 *in planta*. (a) Confocal microscopy images of *N. benthamiana* leaves co-infiltrated with FoTO1-YFP_n_ and either YFP_c_, YFP_c_-FoTO1, CPR-YFP_c_, or calnexin-YFP_c_. Split-YFP signal is pseudocolored in green and chloroplast autofluorescence is shown in red; merge is the overlay of the chloroplast autofluorescence (the scale bar represents 85 μm). Representative images of 3 independent experiments. (b) Quantification of the log2 fold change in taxadien-5α-ol when mCherry (ctrl) or FoTO1-D149A are transiently expressed in *N. benthamiana* leaves along with the indicated TDS and boost enzymes. mCherry (ctrl) is used as a negative control. Values ± SD of three independent experiments (∗p ≤ 0.05, ∗∗p ≤ 0.01, ∗∗∗p ≤ 0.001, ∗∗∗∗p ≤ 0.0001 by Student’s t test and not shown if insignificant (p > 0.05)). (c) Immunoblot analysis of FoTO1, TDS, mCherry, and GFP co-expressed in *N. benthamiana* under differing ODs. FoTO1 and GFP are expressed at a constant OD in all samples. The OD of TDS *Agrobacterium* increases across the samples as indicated while the OD of mCherry *Agrobacterium* decreases concomitantly. The blue shapes represent the OD infiltrated of the indicated proteins and No-Stain^TM^ is used as a loading control. Representative image of 3 independent experiments.

Given FoTO1’s localization near the surface of the ER and the enrichment of plastidial and ER proteins by TurboID-FoTO1, we wondered whether FoTO1’s non-catalytic boost to pathway yield could depend on the subcellular localization of the typically plastidial TDS. When a truncated, cytoplasmic TDS (cytTDS), indicated boost enzymes, and the wildtype ER-localized T5αH were co-infiltrated with the catalytically inactive mutant FoTO1-D149A, no significant change in taxadien-5α-ol levels was observed. This was in stark contrast to the significant 3-fold increase in taxadien-5α-ol observed when the native chloroplast-localized TDS (chlTDS) was used (**Figure 4b**). The yields of the rearrangement products OCT and iso-OCT followed a similar trend, with a significant reduction in the yield boost provided by non-catalytic FoTO1 when TDS was truncated and localized to the cytosol instead of the chloroplast (**Supplementary Fig 14**). Given the enrichment of ER and plastidial proteins observed in the TurboID dataset as well as the major role NTF2 family members play in transport, it is possible that FoTO1, in addition to its catalytic properties, may also facilitate transport of diterpenes from the plastid to the ER.

In addition to a potential role in transport, our co-immunoprecipitation data above led us to hypothesize that FoTO1 may have a role in the physical organization of enzymes in the Taxol pathway, possibly scaffolding a metabolon. Because a metabolon is important when flux is low, we wondered whether we would only see a need for organization under low-flux conditions. To investigate how pathway flux may affect FoTO1, we used reconstitution experiments with lower and higher yields of pathway enzymes by adjusting the OD of agrobacterium infiltration. Here, we assumed that every cell was infiltrated and altering the OD affected the amount of expressed protein in a cell for our given range (Carlson, Rajniak and Sattely, 2023) (**Supplementary Fig 15)**. Interestingly, addition of FoTO1-D149A increased yields of taxadien-5α-ol when a high amount of cytoplasmic TDS (and associated boost enzymes HMGR/GGPPS) was used. This was not the case (ie. no yield increase upon addition of FoTO1-D149A) when a standard amount of cytoplasmic TDS and boost enzymes was used (**Figure 4b, Supplementary Fig 14**). The heightened contribution of FoTO1-D149A when tested at higher ODs of cytoplasmic TDS highlights the importance of the relative stoichiometry of pathway enzymes.

Serendipitously, when analyzing the amounts of proteins produced in each case by immunoblot, we noticed that the addition of TDS seemed to influence FoTO1 protein levels. When the OD of the FoTO1-*Agrobacterium* strain infiltrated was held constant, but the OD of TDS-*Agrobacterium* strain was increased, we observed a marked increase in the amount of FoTO1 but not GFP as measured by immunoblot and fluorescence readout of mTurq-tagged FoTO1 (**Fig 4c, Supplementary Fig 16**). These results indicate that higher ODs of TDS strains substantially elevate FoTO1 protein levels (FoTO1 strain OD is held constant), which may explain why FoTO1-D149A exhibits divergent effects depending on the amount and localization of TDS. We hypothesize that FoTO1 has roles in transport and organization, accounting for the prominence of FoTO1’s non-catalytic contributions when TDS is plastid-localized but not cytosolic. When cytoplasmic TDS is abundant, FoTO1’s enhanced contribution may instead reflect increased FoTO1 protein levels (perhaps through increased stability of FoTO1 through interaction with taxadiene or TDS), which may be required at higher local concentrations in the cytoplasm to execute its organizational effects. These findings suggest that FoTO1’s non-catalytic functions manifest under two distinct conditions, characterized by high cytosolic or low plastidial production of its metabolic intermediate. Together, this observation hints at potential additional mechanisms by which *Taxus* proteins might interact and influence protein stability.

### FoTO1 acts on diverse diterpene pathways

Our data reveal that FoTO1 has both catalytic and organizational functions that contribute to Taxol biosynthesis. Interestingly, FoTO1 seems to co-localize with many P450s, and its impact is influenced by the localization of TDS, suggesting a possible role in the transport of diterpene intermediates from the chloroplast or the organization of pathway enzymes. To investigate whether these organizational roles of FoTO1 are applicable to other pathways, we included FoTO1 in the reconstitution of diterpene pathways from plants unrelated to *Taxus*. Forskolin, a diterpene produced in *Coleus forskohlii*, is a potent adenylate cyclase activator that may have efficacy against HIV^51,52^. The biosynthetic route to forskolin is similar to that of Taxol, with two diterpene synthases (*Cf*TPS2, *Cf*TPS3) in the plastid and a host of subsequent modifying enzymes in the ER^53^. Infiltration of *N. benthamiana* with the early forskolin pathway enzymes *Cf*TPS2, *Cf*TPS3, and *Cf*CYP76AH15 results in the production of the forskolin intermediate 11-oxo-13R-manoyl oxide. Interestingly, when FoTO1 is co-infiltrated with these three forskolin enzymes, there is a 3-fold increase in the production of the forskolin intermediate 11-oxo-13R-manoyl oxide. Addition of the catalytically inactive FoTO1-D149A leads to the same 3-fold increase but addition of a FoTO1 homolog from *A. thaliana* does not (**Figure 5a, Supplementary Fig 15**).

**Figure 5:**
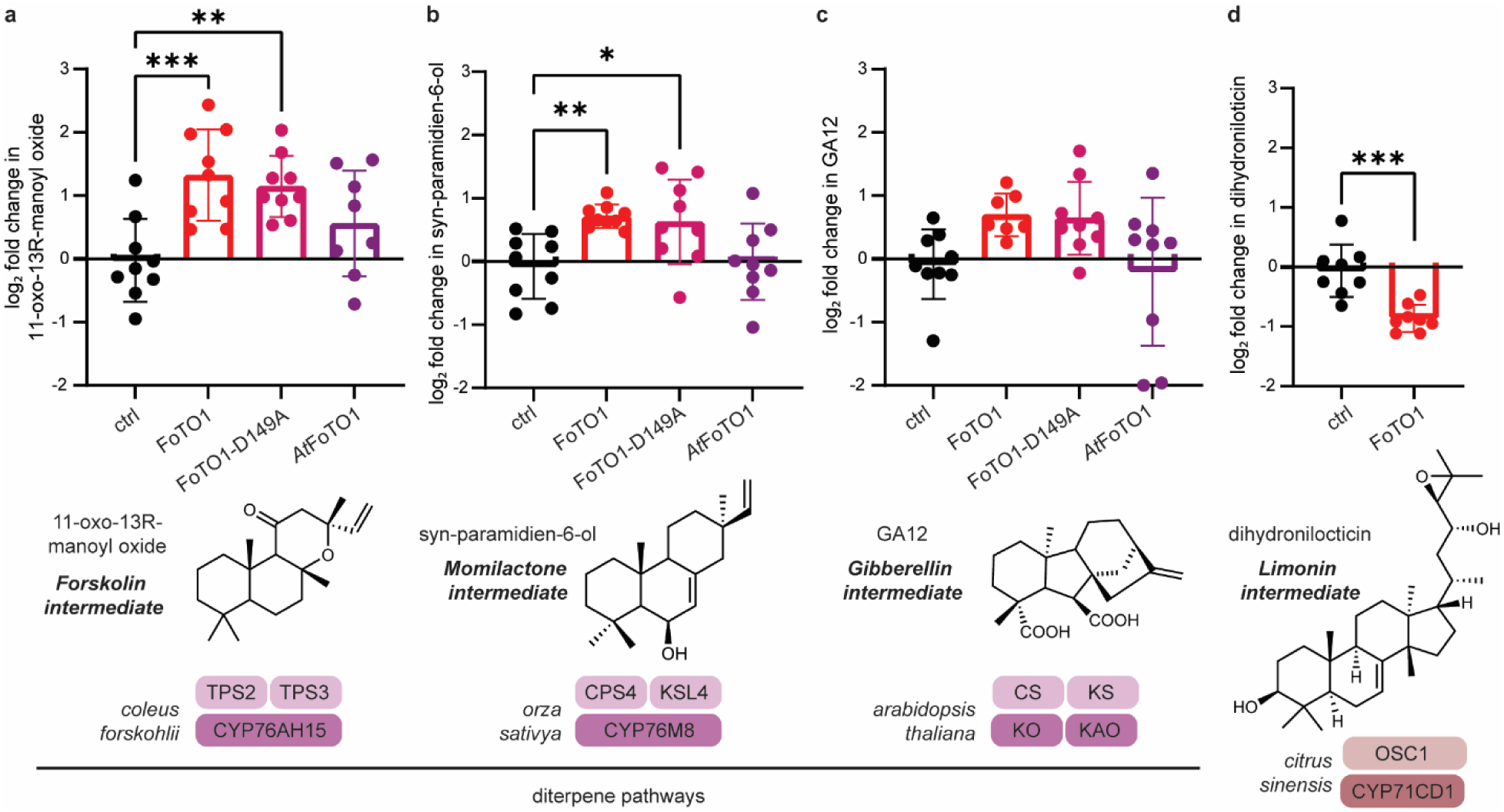
Effect of FoTO1 on diverse terpene pathways from across plant phylogeny. (a) Quantification of the log2 fold change in 11-oxo-13R-manoyl oxide when forskolin early pathway enzymes (HMGR, chlGGPPS, *Fs*TPS2, *Fs*TPS3, *Fs*CYP71AH15) are transiently expressed in *N. benthamiana* leaves with either ctrl (mCherry), FoTO1, FoTO1-D149A, or *At*FoTO. (b) Quantification of the log2 fold change in syn-paramidine-6-ol when momilactone early pathway enzymes (HMGR, chlGGPPS, *Os*KSL4, *Os*CPS4, *Os*CYP76M8) are transiently expressed in *N. benthamiana* leaves with either ctrl (mCherry), FoTO1, FoTO1-D149A, or *At*FoTO. (c) Quantification of the log2 fold change in gibberellin A12 when gibberellin early pathway enzymes (HMGR, chlGGPPS, *At*CS, *At*KS, *At*KO, *At*KAO) are transiently expressed in *N. benthamiana* leaves with either ctrl (mCherry), FoTO1, FoTO1-D149A, or *At*FoTO. (d) Quantification of the log2 fold change in dihydroniloticin when limonin early pathway enzymes (HMGR, cytGGPPS, *Cs*OSC1, *Cs*CYP71CD1) are transiently expressed in *N. benthamiana* leaves either ctrl (mCherry) or FoTO1. mCherry (ctrl) is used instead of FoTO1 as a negative control. Values ± SD of nine independent experiments (∗p ≤ 0.05, ∗∗p ≤ 0.01, ∗∗∗p ≤ 0.001, ∗∗∗∗p ≤ 0.0001 by ordinary one-way ANOVA and not shown if insignificant (p > 0.05)).

We next performed the same experiment on the early steps of the momilactone pathway (from rice) and gibberellin diterpene pathway (from *Arabidopsis* but ubiquitous across all land plants). Neither of these pathways involves resolution of an epoxide. When we included FoTO1 or FoTO1-D149A with the early momilactone and gibberellin pathways enzymes *Os*KSL4, *Os*CPS4, and *Os*CYP76M8 or *At*CS, *At*KS, *At*KO, and *At*KAO respectively, we saw a similar 2 to 3-fold increase in yield (**Figure 5b**)^41,54^. Here, we again saw no significant increase in yield when using a homolog of FoTO1 from *A. thaliana.* Addition of FoTO1 to a triterpene limonoid pathway, however, did not lead to an observed yield increase (**Figure 5c-d**)^55^. This suggests that FoTO1’s non-catalytic functions are relevant to many plastidial diterpene pathways but not cytoplasmic triterpene pathways.

We were also interested in evaluating whether this yield improvement in diterpene pathways was dependent on the localization of the relevant terpene synthases, so we repeated these experiments with truncated terpene synthases that were cytoplasmic and not localized to the plastid. Here, FoTO1 only led to a 1.5-fold yield increase in the cytoplasmic forskolin and momilactone pathways (**Supplementary Fig 17**). This is consistent with our earlier results – in all three diterpene pathways tested, catalytically inactive FoTO1 leads to a 3-fold yield increase which is reduced to a 1.5-fold yield increase when the terpene synthases are rendered cytoplasmic (**Figure 4b, Supplementary Fig 17**). We suspect this may indicate a general role for FoTO1 transporting metabolites between the chloroplast and the ER, perhaps through direct organelle-organelle contacts. In conclusion, these results suggest the non-catalytic roles of FoTO1 to be generally applicable to diterpene biosynthesis in plants.

**Figure 6:**
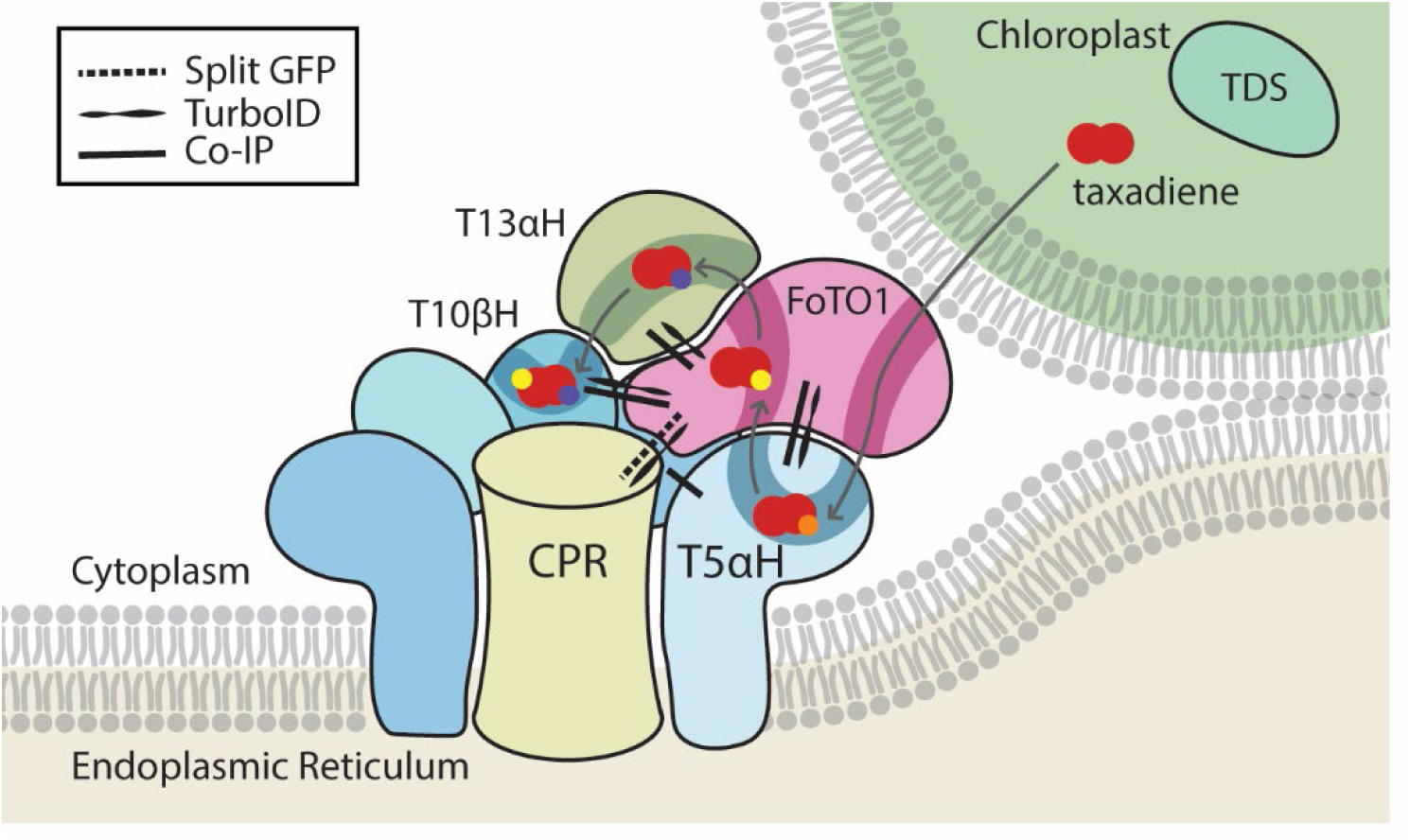
A working model for taxane biosynthesis illustrating multiple independent functions by FoTO1. A working model of biosynthetic protein organization for Taxol biosynthesis. Protein-protein interactions for which there is experimental data are indicated as shown in the legend. Red dots represent the pathway taxanes as they are modified through the pathway

## Discussion

Here, we discover how the recently discovered lynchpin protein FoTO1, which is critical for engineering taxane production in yeast and plant heterologous hosts, streamlines Taxol biosynthesis. Notably, this involves two critical roles, including: (1) catalysis and (2) protein scaffolding, which likely anchors a metabolon for Taxol production. We demonstrate that FoTO1 specifically catalyzes the conversion of the T5αH product taxadiene-4(5)-epoxide to taxadien-5α-ol, resolving a long-standing bottleneck for developing a Taxol bioproduction route. Mutations of the catalytic residues in FoTO1 eliminate its ability to increase the yield of taxadien-5α-ol *in vitro* but only partially reduce this effect *in planta*. FoTO1 increases the yield of taxadien-5α-ol three-fold through non-catalytic mechanisms *in planta* by orchestrating weak protein-protein interactions and increasing proximity between pathway enzymes. These data indicate that enzymatic and scaffolding functions can be decoupled in FoTO1, allowing FoTO1 to enable efficient production of the Taxol intermediate taxadien-5α-ol in heterologous hosts and diverse diterpene pathways.

FoTO1 is a founding member of a small, but increasingly appreciated group of proteins that have clear roles in orchestrating biosynthetic pathways through non-catalytic mechanisms. Two primary types of multifunctionality within this group include enzymes that can act as transporters (such as lipocalin-type prostaglandin D synthase) and those that form metabolons^56–58^. While enzyme complexes have been extensively characterized in primary metabolism (ex. TCA cycle enzymes^59,60^), they are less understood in secondary metabolism in plants. Three lead examples of metabolons in secondary plant metabolism include: GAME15 in steroidal glycoalkaloid biosynthesis, the dhurrin metabolon in cyanogenic glucoside biosynthesis, and complexes in the phenylpropanoid pathway^42,45,46,61^. However, the transient nature of these assemblies makes them challenging to identify and study, making it plausible that metabolons exist in many other biosynthetic pathways and have thus far remained uncharacterized. Consequently, FoTO1 may represent a broader phenomenon in plant biosynthesis that is difficult to observe because of the weak and transient nature of the protein-protein interactions governing this phenomenon, despite their clear importance in driving efficient production in metabolic pathways^62,63^. It is notable that the epoxide intermediate has a relatively short half-life *in planta* compared to other secondary metabolic intermediates. The same is true in the metabolon-containing dhurrin pathway - one of the key intermediates in this pathway is also labile^42^. Thus, close handoff from one protein to the next may facilitate productive metabolism by limiting spontaneous chemistry. These non-catalytic contributions to biosynthesis result in significant increases in the efficiency of these pathways, highlighting the importance of understanding the mechanism of FoTO1.

Unexpectedly, the non-catalytic role of FoTO1 can be used to support other diterpene pathways from outside *Taxus sp*. A similar 3-fold increase in the forskolin, gibberellin, and momilactone pathways is observed upon addition of FoTO1 when these pathways are reconstituted in *N. benthamiana*. This broad functionality suggests that FoTO1 may act as a versatile facilitator of diterpene biosynthesis through non-catalytic mechanisms, potentially by modulating pathway flux, organelle contacts, intermediate transport, or promoting productive protein-protein interactions. These results highlight the importance of the non-catalytic roles of FoTO1 and suggest interactions with conserved proteins (perhaps also present in *N. benthamiana*) or universal features of FoTO1 are critical for its organizational roles in biosynthesis. Understanding the precise mechanisms by which FoTO1 influences diterpene biosynthesis will be critical for enhancing the production of diterpenes in heterologous hosts at scale. Our findings raise questions about the role of FoTO1 homologs across plant species. Nearly every land plant with a sequenced genome has a FoTO1 homolog, and some have several. For instance, the sequenced *Taxus* genome contains three FoTO1 paralogs and 23 NTF2-like proteins, while the *A. thaliana* genome contains 1 FoTO1 paralog and 203 NTF2-like proteins. Despite the prevalence of these FoTO1 paralogs, none outside of *Taxus* have been functionally characterized.

FoTO1 and its homologs belong to the NTF2-superfamily of proteins, a diverse group of proteins that share a conserved NTF2 fold. While FoTO1 and its homologs have not been characterized, various NTF2-like family members have been extensively characterized and the first NTF2 proteins were originally identified for their role in the transport of Ran to the nucleus, which is conserved among its orthologs in plants^25,27,64^. Other plant NTF2-like superfamily members have been shown to mediate response to abiotic stress but none have yet been implicated in biosynthesis^28,65^. A key feature among NTF2-like superfamily proteins is their exceptionally versatile fold which grants them the ability to bind small molecules tightly^25^. The versatility of this fold and its ability to bind small molecules have been exploited by evolution to generate NTF2 superfamily members with roles in both transport and catalysis^50,64,66–68^. More recently, the structural adaptability of this fold has been leveraged in computational approaches to protein design to create novel small-molecule binding proteins, sensors, and enzymes^69–72^. Given the extreme versatility of this fold and its ability to bind small molecules with high affinity, it is likely that many other uncharacterized NTF2-like proteins exist in plants with important roles in specialized metabolism and biosynthesis.

It is notable that some taxadien-5α-ol product is produced spontaneously even in the absence of FoTO1 catalysis. This represents a unique opportunity to decouple scaffolding and catalytic roles in a biosynthetic enzyme. We suspect that other proteins in the pathway are also involved in protein-protein interactions and contribute at some level to pathway organization but the transient nature of the observed protein-protein interactions and the difficulty detecting non-catalytic contributions of biosynthetic enzymes may have limited our understanding of the full extent to which structural and organizational roles contribute to pathway efficiency. Expanding the use of proximity labeling and interaction-based assays may reveal that multifunctionality is a more common feature of plant biosynthetic enzymes than previously recognized. More importantly, these assays may reveal how the spatial arrangement of enzymes is critical for their function. Metabolons and other assemblies of proteins offer cells the opportunity to exploit low or high concentrations of intermediates and to reduce the entropic cost for metabolite channeling. Understanding and harnessing these biosynthetic strategies could provide valuable advantages for further pathway engineering in both plant and non-plant heterologous hosts, allowing for streamlined biomanufacture of clinically relevant biotherapeutics including Taxol.

## Methods

### Cloning

Proteins were tagged for Co-IP, immunoblot, TurboID, and BiFC by amplifying genes from plasmids or *Agrobacterium* glycerol stocks using PCR (PrimeStar, Takara Bio R045B, primers in Supplementary Table 3). The indicated PCR products were then purified using Zymo DNA Clean & Concentrator (Zymo #D4004) and, using HiFi DNA assembly mix (New England Biolabs), ligated into pEAQ-HT vector linearized with AgeI and XhoI (New England Biolabs). The ligation mixture was then transformed into 10-beta competent *E. coli* cells (New England Biolabs). Following overnight incubation of the competent cells on plates, single colonies were picked and grown in liquid media overnight. Plasmid DNA was then purified using a ZymoPURE plasmid miniprep kit (Zymo #D4212) and subjected to whole-plasmid sequencing (Plasmidsaurus). FoTO1 mutant plasmids were created using the same protocol. PCR fragments used to create *E. coli* and yeast expression vectors were ligated into vectors specified in Supplementary Table 3.

### Expression of FoTO1 in yeast

Yeast cultures were grown to saturation in 4 mL YPD medium supplemented with 400 µg mL⁻¹ G418 in glass culture tubes for 2 days. Cells were harvested by centrifugation at 1,000 × g for 5 min, resuspended in YPG induction medium (10 g L⁻¹ yeast extract, 20 g L⁻¹ peptone, 20 g L⁻¹ galactose), and incubated overnight. Cells were subsequently lysed by the addition of 0.5 mm glass beads and 100 µL ethyl acetate, followed by vigorous vortexing. The resultant culture was spun down for 2 minutes at 18000xg, and 50uL of the organic layer was loaded onto the GCMS.

### Purification of FoTO1 from E. Coli

FoTO1 was cloned into the pET28a expression vector and transformed into *E. coli* BL21(DE3) Star competent cells as described previously. Transformants were selected on LB agar plates supplemented with kanamycin (50 µg mL⁻¹). Single colonies were used to inoculate 2 mL LB starter cultures containing kanamycin (50 µg mL⁻¹) and grown overnight, which were then used to seed 200 mL LB cultures supplemented with kanamycin. Cultures were grown at 37 °C to an optical density at 600 nm (OD₆₀₀) of ∼0.6, after which protein expression was induced with 0.2 mM IPTG and cultures were shifted to 18 °C. Cells were harvested after an additional 18 h by centrifugation and pellets were either processed immediately or stored at −80 °C.

Cell pellets of FoTO1 or point mutants were resuspended in 15 mL lysis buffer (20 mM HEPES, 300 mM NaCl, 10 mM imidazole, 1.5% Igepal, pH 8.0) and lysed by three passes through an air-driven high-pressure homogenizer (EmulsiFlex B15, Avestin). Lysates were clarified by centrifugation at 21,000 × g for 25 min at 4 °C. The supernatant was loaded onto Ni–NTA agarose resin (QIAGEN), washed sequentially with 50 mL wash buffer 1 (20 mM HEPES, 300 mM NaCl, 20 mM imidazole, pH 8.0) followed by 15 mL wash buffer 2 (20 mM HEPES, 300 mM NaCl, 50 mM imidazole, pH 8.0), and eluted with elution buffer containing 500 mM imidazole. Eluted fractions were analyzed by SDS–PAGE, pooled as appropriate, and dialyzed overnight. Protein was concentrated using 10 kDa molecular weight cutoff centrifugal filters (Amicon, Sigma) and buffer-exchanged into storage buffer (10 mM HEPES–KOH, 50 mM KCl, 1 mM DTT, 1 mM EDTA, 10% glycerol, pH 8.0). Protein concentration was determined using a Bradford assay and corrected for co-eluting impurities. Purified FoTO1 was snap-frozen in liquid nitrogen and stored at −80 °C.

pYYeD60 plasmids containing T5αH or T13αΗ were transformed into WAT11 yeast strains expressing Arabidopsis CPR11 using lithium acetate, and selected using synthetic-dropout medium plates lacking uracil via growth at 30C for 2 days. Colonies were selected and grown to saturation in a starter culture for 1-2 days at 28C. The starter culture was used to inoculate a 250ml culture, and grown to an OD of 3.0 in synthetic-dropout medium lacking uracil. Subsequently, expression was induced by pelleting cells at 1000xg and then altering the medium to yeast peptone extract containing 2% galactose (10mg/L yeast extract, 20mg/L peptone, 20mg/L galactose) for another 16 hours after which this culture was immediately used for microsomal protein isolation, performed as previously described. Microsomal protein was stored in TEG buffer (50 mM Tris-HCl, 1 mM EDTA, 20% [v/v] glycerol, pH 7.4), aliquoted into 1.5 mL microfuge tubes, snap frozen in liquid nitrogen, and stored at −80 °C. Typical yields were around 20-30mg/ml of crude protein, as estimated via the Bradford assay.

### Purification and chemical synthesis of taxadiene and taxadiene epoxide

Taxadiene was purified as we previously described, from the JBEI-18127 strain^17^. Glycerol stocks were used to inoculate 20 mL starter cultures in yeast peptone dextrose (YPD) medium and grown at 30 °C for 48 h. Starter cultures were subsequently used to inoculate 2 L YPD cultures, which were grown at 30 °C until reaching an optical density (OD₆₀₀) of approximately 1. At this point, the incubation temperature was reduced to 20 °C to minimize non-enzymatic hydrolysis of geranylgeranyl pyrophosphate, and cultures were grown for an additional 4 days. Taxadiene production was monitored by small-scale liquid–liquid extraction of culture media followed by GC–MS analysis.

Cultures were harvested by centrifugation at 3,000 × g to separate cells from media, and each fraction was extracted independently. Cell pellets were frozen at −80 °C for 1 h to facilitate lysis, then incubated overnight with three volumes of ethyl acetate. Culture supernatants were subjected to three successive liquid–liquid extractions with ethyl acetate. Organic extracts from both fractions were combined and concentrated by rotary evaporation. The resulting crude extract was weighed, redissolved in a minimal volume of hexane, and loaded onto a silica gel column at a 1:40 (w/w) product-to-silica ratio for flash chromatography. Isocratic elution was conducted using hexane as the mobile phase, collected in 5ml fractions, which were analyzed by GC–MS to identify taxadiene-containing fractions. Fractions were subsequently concentrated via rotary evaporation, with purity confirmed via NMR. Typical production yields were approximately 10–15 mg L⁻¹, with isotaxadiene present at an approximate 1:10 ratio relative to taxadiene, as monitored via NMR.

The chemical synthesis of taxadiene 4(5)-epoxide from taxadiene was performed as previously described by Barton *et al.* Dimethyldioxirane was generated *in situ* by addition of oxone (5 g) to a chilled solution containing H₂O (4 mL), acetone (6 mL), and NaHCO₃ (4.8 g), followed by stirring at 4 °C for 20 min^35^. DMDO was subsequently isolated by collection in a bump trap during rotary evaporation of the reaction mixture at 155 mmHg, with the bath temperature ramped from room temperature to 40 °C over 15 min. The concentration of DMDO was determined by 3 iodometric titrations using the method described by Ahmat, et al, to be 13.25mM. Taxadiene (2.44 mg, 8.95 μmol) was dissolved in dichloromethane (150 μL) and stirred at 4 °C for 5 min. 0.7 molar equivalents of DMDO (50 μL, 6.27 μmol) was added dropwise, after which the reaction mixture was allowed to warm to ambient temperature and stirred for 30 min^73^. Reaction progress and conversion of the starting material were monitored by GC–MS analysis. Reaction products were isolated by rotary evaporation and were not subjected to further purification due to the previously reported instability of the epoxide on silica gel.

### In-vitro enzymatic assays

Enzymatic oxidation assays of taxadiene were performed in 100 µL reaction volumes in 1.5 mL microcentrifuge tubes (Eppendorf). Each reaction contained 36 µg purified cytochrome P450 enzyme, 50 µM taxadiene, and 20 mM HEPES buffer supplemented with 10 mM NaCl (pH 7.5). Reactions were incubated for 3 h at 30 °C with shaking at 250 rpm in a thermal shaker. Following incubation, reactions were extracted with 120 µL ethyl acetate. For reactions involving FoTO1, 2.8nM FoTO1 was added after completion of the 3 h P450 reaction and incubated for an additional 2 h. Samples were vortexed vigorously and centrifuged at 18,000 × g to separate the organic and aqueous phases. A 50 µL aliquot of the organic phase was analyzed by GC–MS.

Enzymatic assays examining the direct reaction of FoTO1 with taxadiene 4(5)-epoxide were conducted in liposome preparations due to substantially higher rate of FoTO1 catalysis (**Supplementary Fig 3**). Liposomes were prepared from phosphatidylcholine (soy; Cayman) at a final concentration of 2 mg L⁻¹ in 10 mM HEPES buffer containing 20 mM sodium cholate. The mixture was sonicated for 8 min using a probe-tip sonicator with 2 s on/off cycles at an amplitude of 20. Liposome stocks were allotted and stored at −20 °C until use. Reactions were performed with 1.4 nM FoTO1 and 25 µM substrate in 20 mM HEPES buffer containing 10 mM NaCl (pH 7.5).

For kinetic analysis and determination of K_m_ values, reactions were conducted in duplicate using varying concentrations of taxadiene 4(5)-epoxide (4–200 µM) and 10 µM linalool as an internal standard. Multiple 50 µL reaction aliquots were prepared and quenched at short time points (30 s, 1 min, 2 min, and 3 min) corresponding to the linear phase of product formation. Reactions were quenched by flash freezing in liquid nitrogen and extracted three times with 100 µL ethyl acetate. Organic extracts were combined and concentrated under a gentle stream of air and redissolved in 50 µL ethyl acetate. Conversion of substrate to product was quantified by GC–MS by summing all degradation peaks corresponding to the starting material and calculating the apparent increase in the taxadien-5α-ol product peak, normalized to linalool recovery, whose average was used to plot the rate of reaction.

### Transient expression in *Nicotiana Benthamiana* via infiltration with agrobacterium

The pEAQ-HT plasmids harboring the relevant genes were introduced into *Agrobacterium tumefaciens* (strain GV3101) cells via the freeze–thaw transformation method. The transformed cells were plated on 523-agar (Phytotech Labs) medium supplemented with kanamycin (50 μg/ml) and gentamicin (30 μg/ml) and incubated at 30°C for 2 days. Single colonies were then picked and cultured overnight at 30°C in liquid 523 medium containing both antibiotics. The overnight cultures were used to prepare 25% glycerol stocks, which were stored at −80°C for long-term preservation.

For subsequent *Nicotiana benthamiana* infiltration experiments, Agrobacterium glycerol stocks were streaked onto 523-agar plates with kanamycin and gentamicin and incubated for 1–2 days at 30°C. Cells were scraped off using a 10-μl inoculation loop and resuspended in 1 ml of Agrobacterium induction buffer (10 mM MES, pH 5.6; 10 mM MgCl₂; 150 μM acetosyringone; Acros Organics) in 2-ml safe-lock tubes (Eppendorf). The suspensions were vortexed briefly to ensure homogeneity, followed by a 2-hour incubation at room temperature. OD600 readings were taken for each suspension, and the final infiltration solution was prepared by adjusting the OD600 to 0.2 for each strain (unless otherwise noted) by diluting with the induction buffer. For infiltration, four-week-old *N. benthamiana* leaves were targeted, using needleless 1-ml syringes to infiltrate the abaxial side. Three to nine biological replicates were performed (as indicated), each corresponding to one leaf per plant.

### Extraction of metabolites in *Nicotiana Benthamiana*

Four days post-*Agrobacterium* infiltration, *N. benthamiana* leaf samples were harvested using a 1 cm diameter leaf disc cutter. Each biological replicate comprised six discs (approx. 60 mg) collected from a single leaf and stored in a 2-ml safe-lock tube (Eppendorf). Samples were immediately flash-frozen and lyophilized overnight. Analyses of the early taxanes taxadiene, taxadien-5α-ol, relevant rearrangements products (iso-OCT, OCT, 2d), 11-oxo-13R-manoyl oxide, syn-paramidien-6-ol, and dihydronilocticin were done by GC–MS, and analyses of GA12 were done using liquid chromatography–mass spectrometry (LC–MS). For metabolite extraction, one 5-mm stainless steel bead and an appropriate solvent - either 750 uL ethyl acetate (ACS reagent grade; J.T. Baker) for GC-MS or 30 uL/mg 80% methanol (high-performance liquid chromatography (HPLC) grade; Fisher Chemical) / 20% water (for LC-MS analysis) - was added to each tube. The tissues were homogenized using a ball mill (Retsch MM 400) at 25 Hz for 2 min and centrifuged at 18,200g for 10 min. GC-MS bound supernatants were transferred to 50-μl glass inserts and placed in 2 ml vials before analysis by the GC-MS instrument. LC–MS samples were filtered using 96-well hydrophilic PTFE filters with a pore size of 0.45 μm (Millipore) and analysed by the LC–MS instrument.

### GC-MS analysis

GC–MS analysis was performed using an Agilent 7820A GC system coupled to an Agilent 5977B single quadrupole mass spectrometer. Data acquisition and processing were conducted via Agilent Enhanced MassHunter and MassHunter Qualitative Analysis (B.07.00), respectively. Compounds were separated on an Agilent VF-5HT column (30 m × 0.25 mm × 0.1 μm) using helium as the carrier gas at a constant flow of 1 ml/min. A 1-μl sample was injected in split mode (10:1 ratio) with an inlet temperature of 280°C. The oven program began at 130°C (2-min hold), increased to 250°C at 8°C/min, and finally reached 310°C at 60°C/min, followed by a 1-min post-run at 320°C. MS detection was initiated after a 4-min solvent delay, covering a mass range of 50–550 m/z at a scan speed of 1,562 u/s. Interface temperatures were maintained at 250°C (transfer line), 230°C (source), and 150°C (quadrupole). Dihydroniloticin analysis followed the protocol established by Hodgson et al ^55^.

### LC-MS analysis of metabolites

Samples of GA12 were analyzed using an Agilent 1260 HPLC system coupled to an Agilent 6520 Q-TOF mass spectrometer, with data acquisition and processing performed via MassHunter Workstation and Qualitative Analysis 10.0, respectively. Chromatographic separation was achieved on a Gemini 5-μm NX-C18 110-Å column (2 x 100 mm; Phenomenex) at a flow rate of 400 μl/min under ambient temperature. The mobile phase consisted of 0.1% formic acid in water (A) and 0.1% formic acid in acetonitrile (B). Following a 2 μl injection, the gradient profile was as follows: an initial 3-min hold at 30% B, a linear ramp to 97% B from 3 to 8 min, a 3-min hold at 97% B, and a final return to 30% B between 11 and 14 min. Mass spectra were acquired in negative electrospray ionization (ESI) mode over a range of 50–1,200 m/z at a scan rate of one spectrum per second. For the 6520 system, source parameters were set to a gas temperature of 325 °C, drying gas flow of 10 l/min, nebulizer pressure of 35 psi, VCap of 3,500 V, fragmentor at 150 V, skimmer at 65 V, and octupole 1 RF Vpp at 750 V. MS/MS fragmentation was conducted using the [M+Na]+ adduct as the precursor ion with a collision energy of 30 eV, unless otherwise specified.

### Co-immunoprecipitation

Four days post-infiltration, *Nicotiana benthamiana* leaves were harvested and homogenized in liquid nitrogen. The resulting powder was resuspended in extraction buffer (50 mM Tris pH 7.5, 150 mM NaCl, 0.6% NP-40, 0.6% CHAPS, and 1 mM β-mercaptoethanol). Lysates were incubated on ice and clarified by centrifugation at 20,000g (10 min, 4°C), with total protein concentration determined via Bradford assay (Abcam). For immunoprecipitation, 15 μl of protein-G-coated magnetic beads (Invitrogen) were pre-washed in binding buffer (50 mM Na2HPO4, 25 mM citric acid, pH 5.0) and incubated with 1 μl of anti-V5 antibody (Invitrogen #MA1-34099), 1 μl of anti-GFP antibody (ProteinTech #66002-1-Ig), or 1 μl of anti-Myc antibody (ProteinTech #16286-1-AP) for 1 h at room temperature under agitation. Following two washes in extraction buffer, the antibody-bound beads were incubated with 100 μg of total protein lysate (∼40 μl) for 15 min at room temperature. Finally, the bead complexes were washed three times in extraction buffer and prepared for immunoblotting using 4x NuPAGE LDS sample buffer (Invitrogen).

### Immunoblotting

Protein lysates were resolved on a NuPAGE gel (Invitrogen) at 100V for 1.5 h. Proteins were then transferred onto a PVDF membrane using a Bio-Rad Trans-Blot Turbo Transfer System. Membranes were blocked and incubated with primary antibodies (anti-V5 at 1:1,000 (Invitrogen #MA1-34099)), anti-HA-HRP at 1:2,500 (R&D Systems #HAM0601), anti-myc at 1:2000 (ProteinTech #16286-1-AP), anti-His at 1:1000 (Cell Signaling #9991S), anti-GFP at 1:2000 (ProteinTech #66002-1-Ig)) for either 3 h at room temperature or overnight with constant agitation. After washing, the blots were incubated with HRP-conjugated protein G (Genscript; 1:5,000) for 1 h. Protein bands were visualized using an iBright FL1500 Imaging System.

### Bi-molecular fluorescence complementation

Leaves were co-infiltrated with *Agrobacterium* strains at an OD of 0.2 as described above. Four days post infiltration, leaf discs 3/16” in diameter were cut out using leaf disc cutters and wet-mounted onto glass slides by placing them into water-filled wells created using Carolina observation gel (Carolina #132700). Slides were then imaged using a Leica Stellaris 5 microscope. Z-stack images were acquired using the LasX software with a thickness of 10 uM over 30 steps and a maximum intensity projection was created using Fiji (ImageJ).

### Proximity labeling of proteins by TurboID

Leaves were infiltrated with TurboID constructs and indicated proteins at an OD of 0.2. Three days post infiltration, leaves were collected and infiltrated with 30 uM biotin (Thermo Fisher #A14207.03) for 1 hour at room temperature. Leaves were then homogenized for 2 minutes at 25 Hz in liquid nitrogen with steel balls using a ball mill (Retsch MM 400). Following homogenization, leaves were resuspended in 2 mL extraction buffer (50 mM Tris pH 7.5, 150 mM NaCl, 0.1% SDS, 1% Triton-X-100, 0.5% Na-deoxycholate, 1 mM EGTA, 1 mM DTT, 1 mM PMSF, 1x protease inhibitor (Roche #11697498001)), and incubated under agitation for 10 minutes and 4 °C. Lysates were then sonicated on ice (20% amplitude, 10 seconds on / 10 seconds off for 2 minutes (QSonica #Q700)) before centrifugation at 2500 rpm for 5 min at 4 °C. Lysates were desalted using a PD-10 column (Cytiva) according to manufacturer’s instructions. The lysates were subjected to a Bradford assay and equivalent amounts of total protein were mixed with Streptavidin coated magnetic beads and incubated at 4 °C overnight. The following methods are as previously described but briefly summarized below ^47^.

Beads were centrifuged for 2 minutes at 1500 rpm at 4 °C. The streptavidin beads were then washed for 8 minutes twice with 1 mL of cold extraction buffer, twice with cold equilibration buffer (50 mM Tris pH 7.5, 150 mM NaCl, 0.1% SDS, 1% Triton-X-100, 0.5% Na-deoxycholate, 1 mM EGTA, 1 mM DTT), once with cold 1 M KCl, and twice with cold extraction buffer. Beads were transferred to a new tube following the first wash and the KCl wash. Beads were then washed six times for 8 minutes with room temperature MS grade PBS, with transfer to a new tube after the first and sixth wash. The supernatant was removed and beads were stored at −80 °C. The follow up on beads digestion and peptide desalting were described as ^74^.

### LC-MS/MS analysis of proteomic samples

LC-MS/MS was carried out on an Orbitrap Eclipse mass spectrometer (Thermo Fisher), equipped with an Easy LC 1200 UPLC liquid chromatography system (Thermo Fisher). Peptides were first trapped using a trapping column (Acclaim PepMap 100 C18 HPLC, 75 μm particle size, 2 cm bed length), then separated using analytical column AUR3-25075C18, 25CM Aurora Series Emitter Column (25 cm x 75 µm, 1.7 µm C18) (IonOpticks). The flow rate was 300 nL/min. Peptides were eluted by a gradient from 3 to 28 % solvent B (80 % acetonitrile, 0.1 % formic acid) over 106 min and from 28 to 44 % solvent B over 15 min, followed by a short wash (15 min) at 90 % solvent B. Precursor scan was from mass-to-charge ratio (m/z) 375 to 1600 (resolution 120,000; AGC 200,000, maximum injection time 50ms, Normalized AGC target 50%, RF lens(%) 30) and the most intense multiply charged precursors were selected for fragmentation (resolution 15,000, AGC 5E4, maximum injection time 22ms, isolation window 1.4 m/z, normalized AGC target 100%, include charge state=2-8, cycle time 3 s). Peptides were fragmented with higher-energy collision dissociation (HCD) with normalized collision energy (NCE) 27. Dynamic exclusion was enabled for 30s.

### MS data analysis and Label-free quantification

MS/MS spectra were searched against the Uniprotkb_proteome_UP000084051 database (73,700 entries) supplemented with 24 *Taxus* protein sequences using the MSFragger 2.3.0 software with default parameters. Label free quantitative (LFQ) intensities were obtained from the search results. Downstream statistics were analyzed in Perseus (version 2.0.10.0). MaxLFQ intensities were log2 transformed prior to statistical analysis. Proteins were retained if they contained at least two valid values in at least one experimental group (FoTO-TurboID or TurboID). Missing values were imputed from a normal distribution (width=0.3, downshift=1.8) applied column wise. Differential abundance was assessed using a two-sample t-test with permutation-based false discovery rate (FDR) control (250 permutations, FDR = 0.1) and an S0 parameter of 1.

## Acknowledgements

We thank all E.S.S. laboratory members for constructive feedback on this project; D. Wengier for discussions and comments on the manuscript, and C. Liou and S. Niehs for discussions, and M. Brockley for the ER-GFP plasmid. We also thank the Brophy Lab (Stanford University) for the use of their Leica Stellaris 5 microscope, and the Keasling lab (University of California, Berkeley) for the TDS and TDS/T5αH engineered yeast strains used in this study. This work is supported by NIH R01 AT010593 (E.S.S.), a Doerr Accelerator grant (E.S.S.), National Science Foundation Graduate Research Fellowship under Grant No. DGE-2146755 (C.W.), NIH Biotechnology Training Grant 5T32GM141819 (C.W.), NIH K99 1K99AT012787 (C.J.M.), the Damon Runyon Cancer Research Foundation (DRG-2421-21; C.J.M.), NIH S10OD030441 to S.-L.X. and by the Carnegie Endowment Fund to the Carnegie Mass Spectrometry Facility.

## Contributions

C.W., C.J.M., J.C.T.L., and E.S.S. contributed to conceptualization. C.W., A.S., S.K., and S.X. developed the methodology and performed the formal analysis. C.W., A.S., J.C.T.L., and C.J.M. carried out the experiments. C.W. and A.S. wrote the original draft of the manuscript with edits from C.J.M., J.C.T.L., and E.S.S. C.J.M., P.M.F., E.S.S. supervised the project.

