## Supplementary Information for "FoTO1 orchestrates Taxol biosynthesis through catalytic and non-catalytic mechanisms"

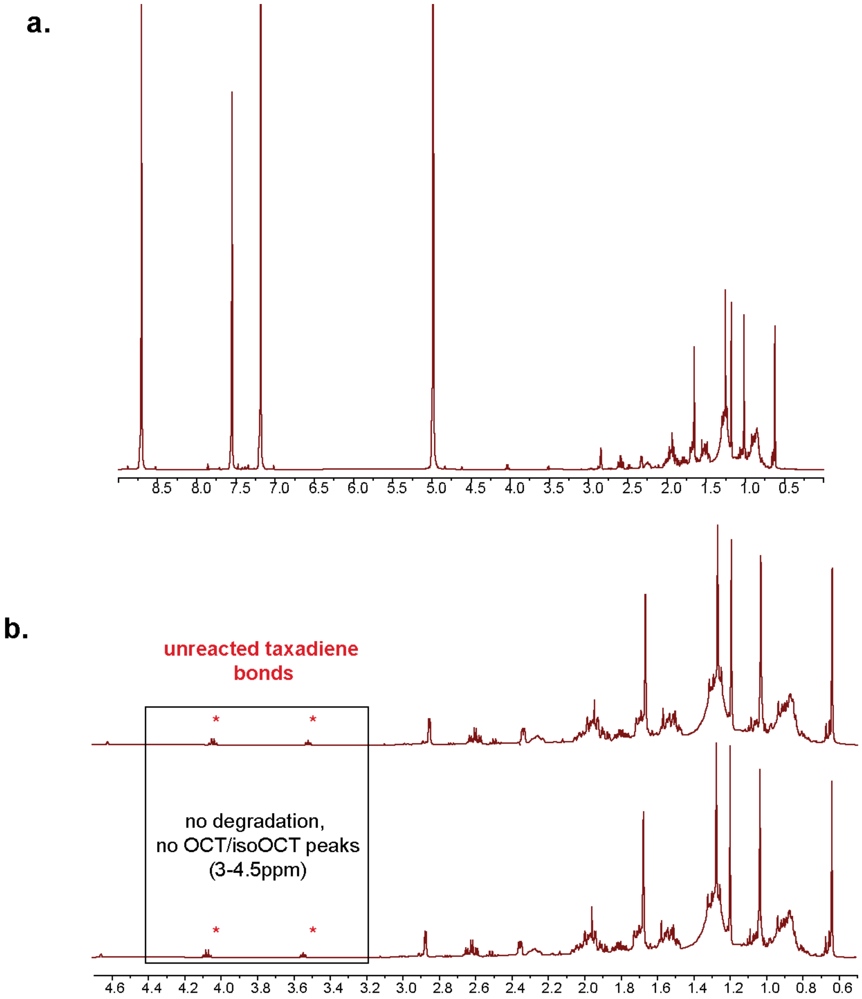


**Supplementary Figure 1: NMR of Taxadiene 4(5)-epoxide sample directly after synthesis and 3 days post synthesis**

**(a)** Full $^{1}H$ NMR (600 MHz) spectrum of crude extract of crude extract producing taxadiene-4(5)-epoxide in $CDCl_3$ (0-9ppm)

**(b)** Comparison of $^{1}H$ NMR spectra after incubation for 3 days at room temperature in $CDCl_3$ to before incubation to monitor reductions in peak height or emergence of novel peaks corresponding to instability.

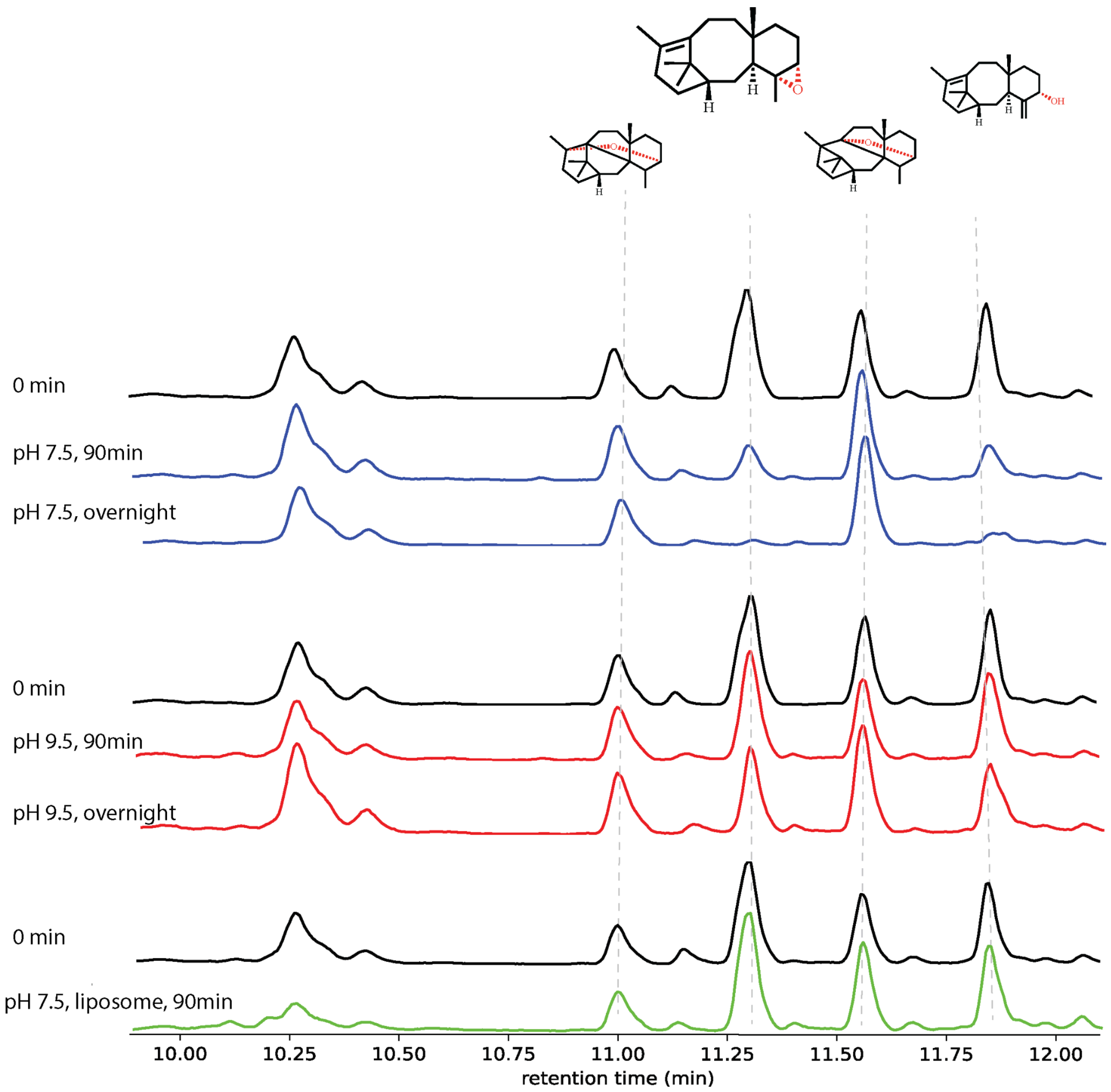


**Supplementary Figure 2: Stability of taxadiene epoxide at different pHs and in the presence of liposomes.**

(**a**) GCMS total ion chromatograms (TICs) of the synthesized taxadiene epoxide directly after synthesis (0 min), after incubation for 90 minutes or after incubation overnight at the indicated conditions. Representative TICs from three independent experiments.

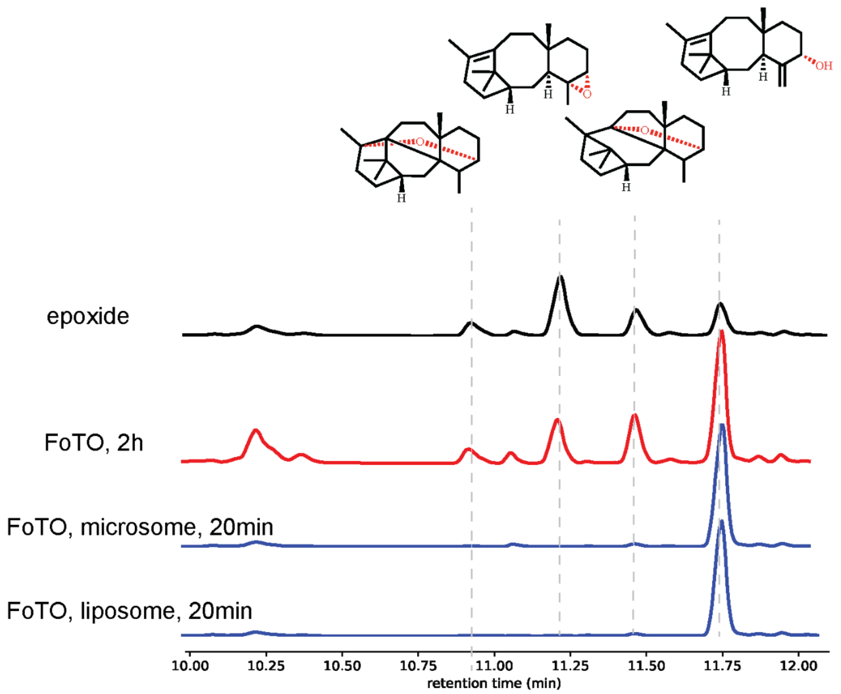


**Supplementary Figure 3: Catalytic activity of FoTO1 increases in the presence of liposomes**.

(**a**) GCMS total ion chromatograms (TICs) *in vitro* incubations of chemically synthesized taxadiene epoxide (**2c**) with 5 nM FoTO1 in solution or in microsomal or liposomal environments (as indicated). Representative TICs from three independent experiments.

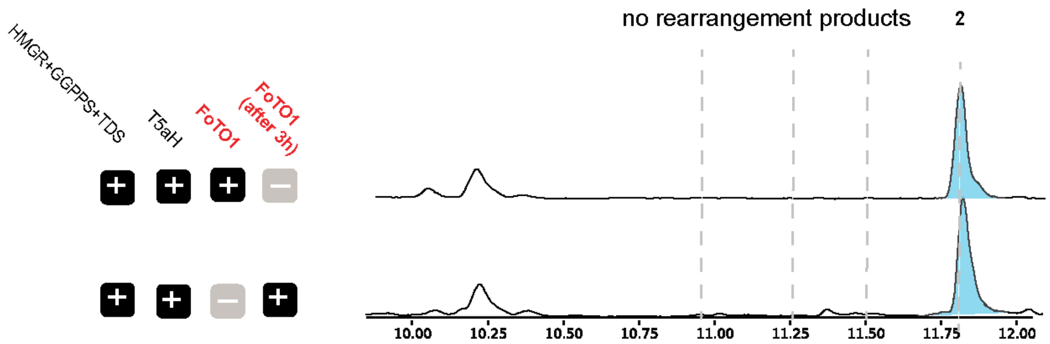


**Supplementary Figure 4: FoTO1 causes major apparent selectivity for taxadiene 5α-ol even when added only after completion of first oxidation.**

(**a**) GC-MS total ion chromatograms of *in vitro* reactions involving taxadiene 5α-hydroxylase co-incubated with purified FoTO1 added directly to the reaction mix or or incubation after completion of reaction (3 h) followed by 1.5h with FoTO1. Representative TICs from three independent experiments.

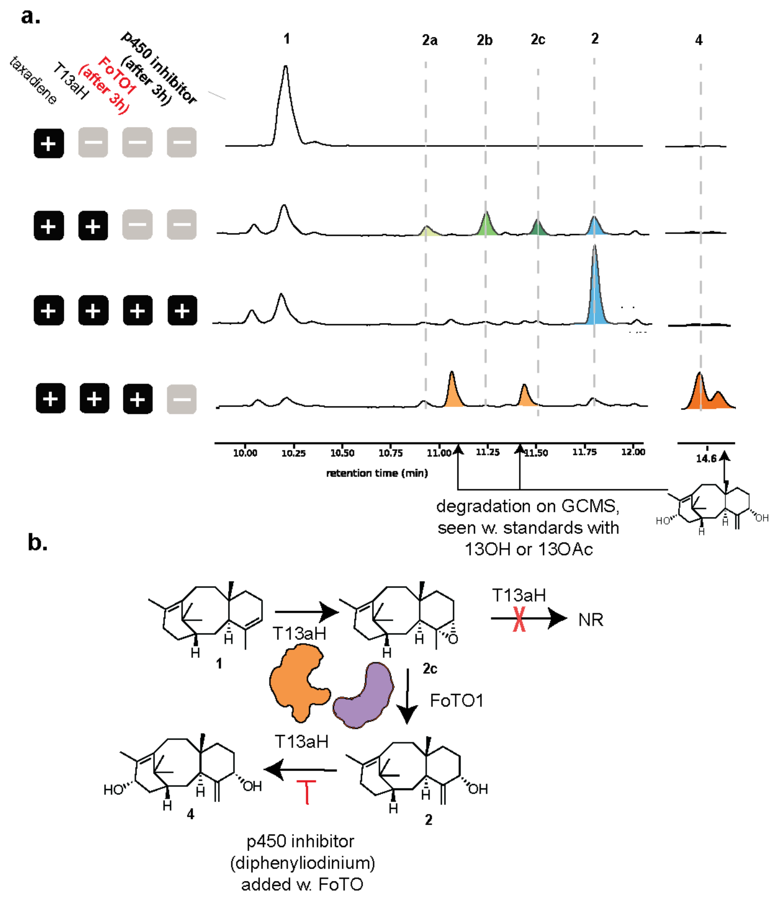


**Supplementary Figure 5: T13αH mechanism occurs analogously to T5αH alongside FOTO**

(**a**) GCMS TICs of T13αH-expressing yeast microsomes, with or without subsequent incubation with FoTO1. Diphenyliodinium was utilized as a p450 inhibitor after addition of FoTO1 to prevent further catalysis by T13αH. Representative TICs from three independent experiments.

(**b**) Schematic of reaction involving T13αH and FoTO1 catalysis of taxadiene.

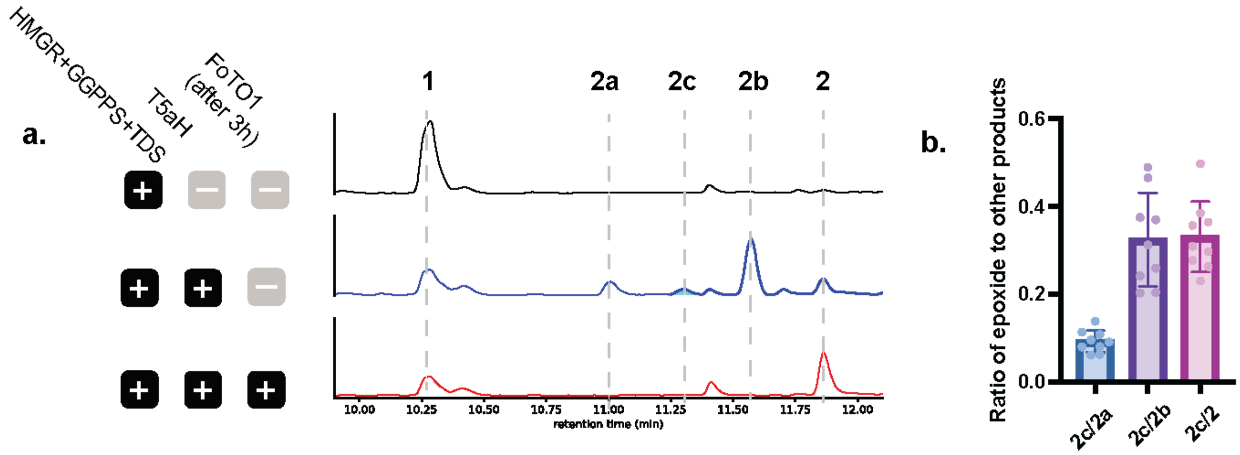


**Supplementary Figure 6: Detection of taxadiene-4(5)-epoxide in *N. benthamiana***

(**a**) GCMS TICs of lysates from *N. benthamiana* infiltrated with low titers of taxadiene 5α-hydroxylase alongside normal titers of taxadiene synthase and diterpene boost genes (HMGR, cytGGPPS). Representative TICs from three independent experiments.

(**b**) Calculated ratio of apparent production of taxadiene-4(5)-epoxide (**2c**) to other degradation products produced in *N. benthamiana* experiments when taxadiene 5α-hydroxylase is expressed at low titers alongside taxadiene synthase and diterpene boost genes.

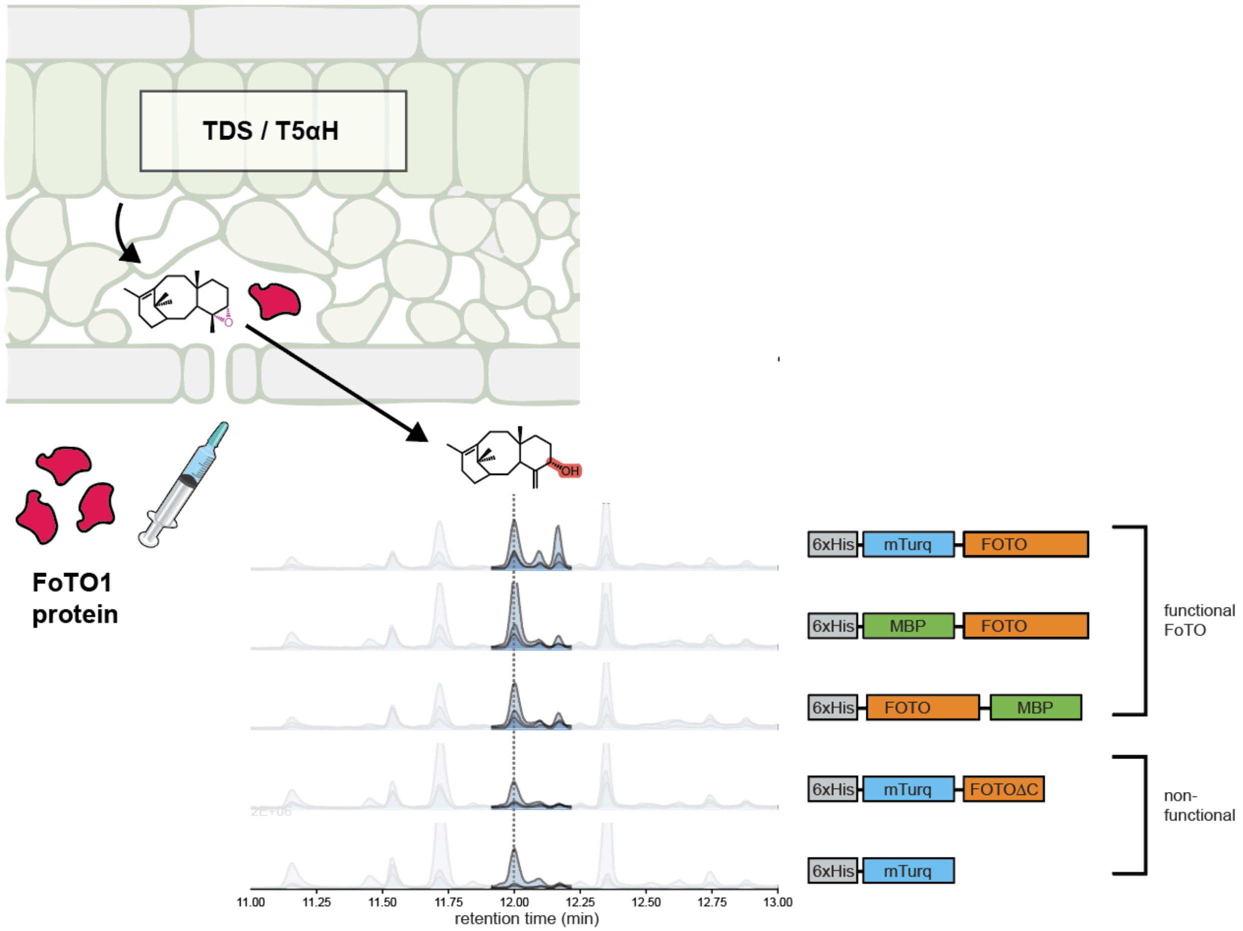


**Supplemental Figure 7**: **FoTO1 addition into leaf leads to increase production of taxadiene 4(5) epoxide**

(**a**) GC-MS total ion chromatograms of taxadiene-5α-ol after purified FoTO1 is incubated with *N. benthamiana* leaf crude extract transiently expressing HMGR, GGPPS, TDS, and T5aH. Representative TICs from three independent experiments.

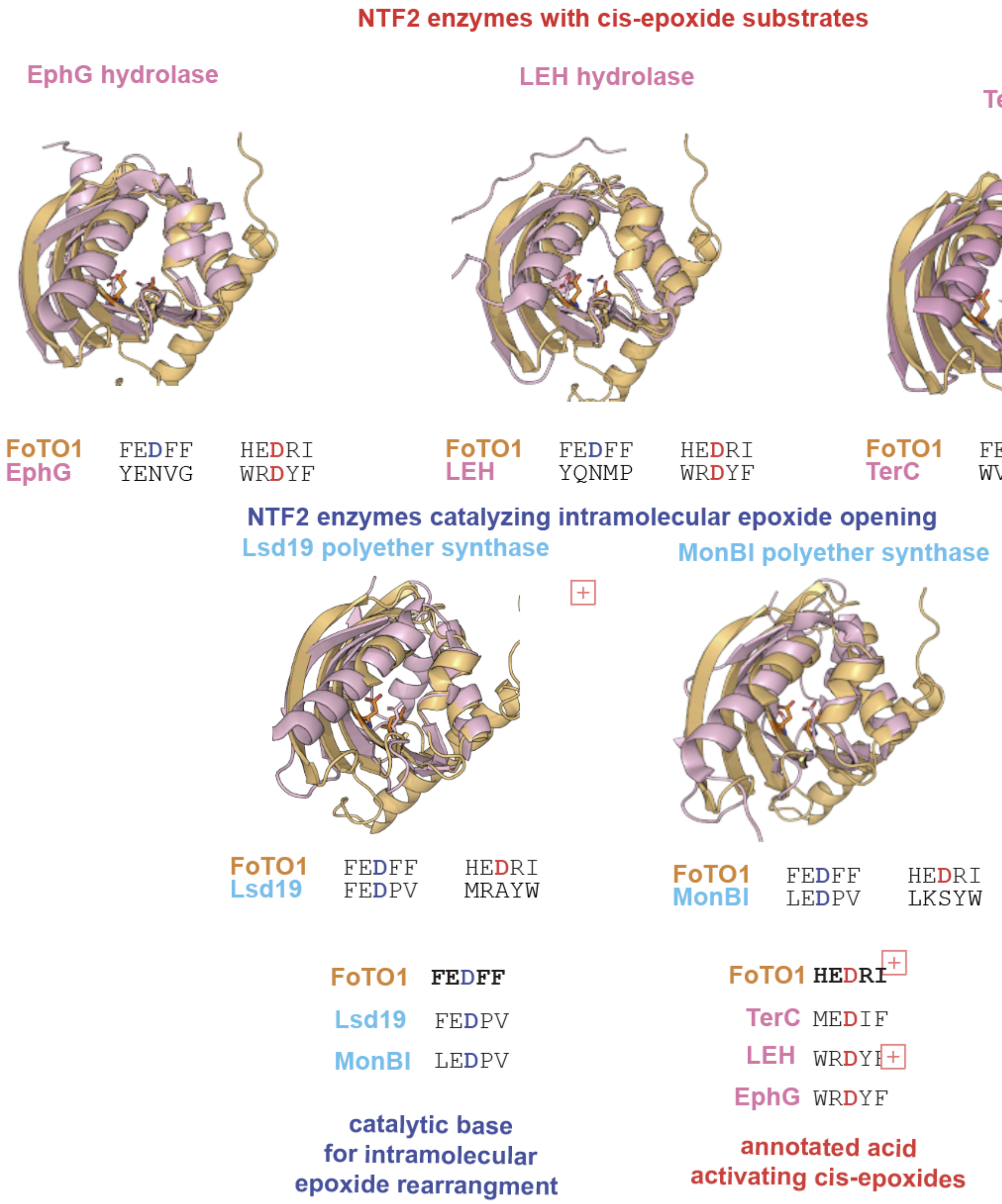


**Supplementary Fig 8: FoTO1 is structurally similar to five epoxide hydrolases and they share spatially conserved residues**

(**a**) Pairwise alignment of FoTO1 with NTF2 enzymes utilizing epoxide substrate or intermediates, with comparisons upon enzymes which either act on similar cis-epoxides or catalyze intramolecular rearrangements. The amino acid sequences that align to D68 and D149 are pictured below the structures.

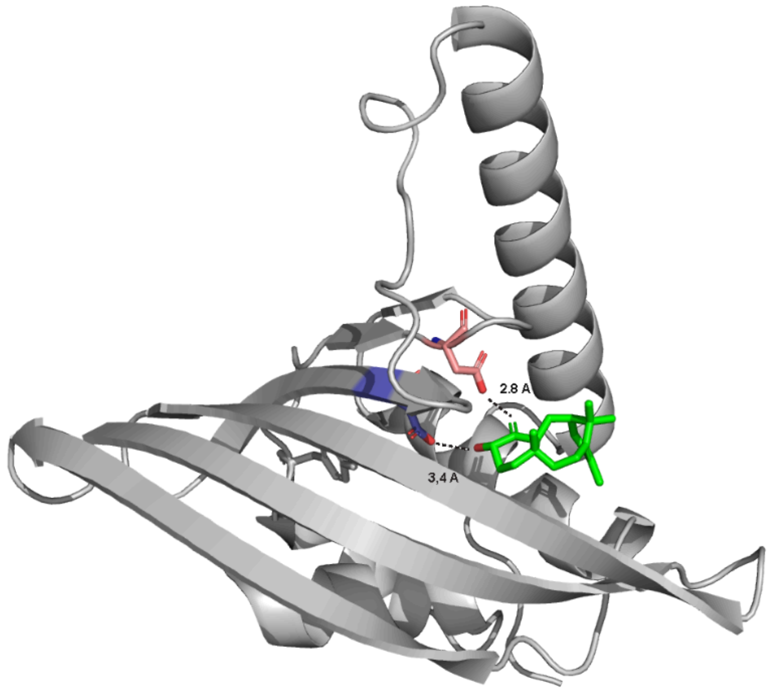


**Supplemental Figure 9: Taxadien-5a-ol computationally docked into the predicted structure of FoTO1**

(**a**) Docking of taxadien-5a-ol into the AlphaFold predicted structure of FoTO1 using DiffDock. The distance between the residues D149 (blue) and D68 (red) in FoTO1 and the protons transferred during epoxide opening and allylic alcohol formation from taxadien-5a-ol are shown as dotted lines.

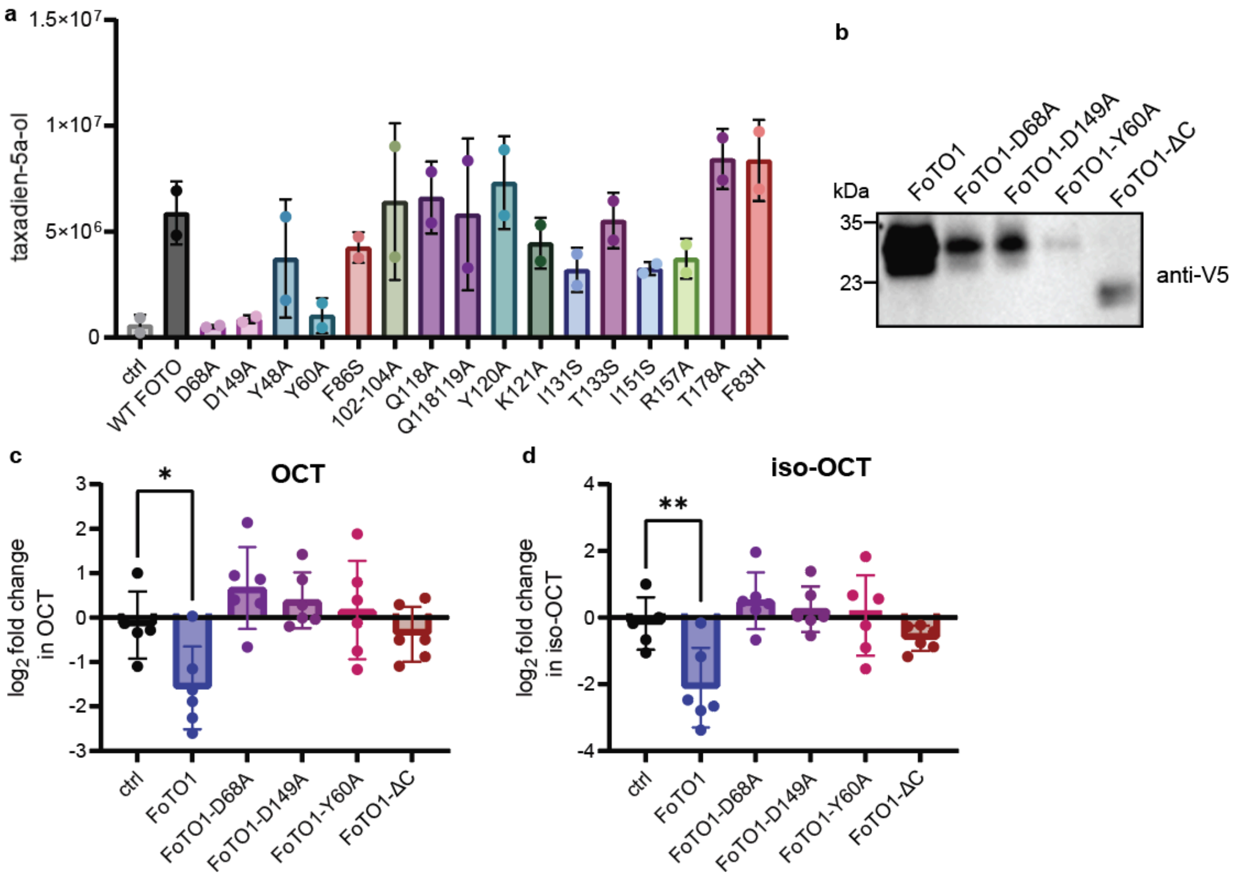


**Supplementary Fig 10: FoTO1 residues D68, D149, and Y60 are crucial for epoxide turnover**

(**a**) Bar graph showing integrated peak area of taxadien-5α-ol when point mutants of FoTO1 are transiently expressed in *N. benthamiana* leaves along with boost and early pathway enzymes (HMGR, GGPPS, TDS1, TDS2, T5αH). Ctrl, mCherry is used instead of FoTO1 as a negative control and wild-type FoTO1 is used as a positive control. Fold change is calculated as the ratio of the measured peak area of taxadien-5α-ol in a sample divided by the average measured peak area of taxadien-5α-ol in the absence of FoTO1 (control) of that run. Values ± SD of two independent experiments (∗$p \leq 0.05$, ∗∗$p \leq 0.01$ by Student's t test and not shown if insignificant ($p > 0.05$)).

(**b**) Immunoblot analysis of indicated V5-tagged FoTO1 mutants.

(**c-d**) Bar graph showing integrated peak area of (**c**) OCT and (**d**) iso-OCT when point mutants of FoTO1 are transiently expressed in *N. benthamiana* leaves along with boost and early pathway enzymes (HMGR, GGPPS, TDS1, TDS2, T5αH). Ctrl, mCherry is used instead of FoTO1 as a negative control and wild-type FoTO1 is used as a positive control. Fold change is calculated as the ratio of the measured peak area of taxadien-5α-ol in a sample divided by the average measured peak area of taxadien-5α-ol in the absence of FoTO1 (control) of that run. Values ± SD of six independent experiments (∗$p \leq 0.05$, ∗∗$p \leq 0.01$ by Student's t test and not shown if insignificant ($p > 0.05$)).

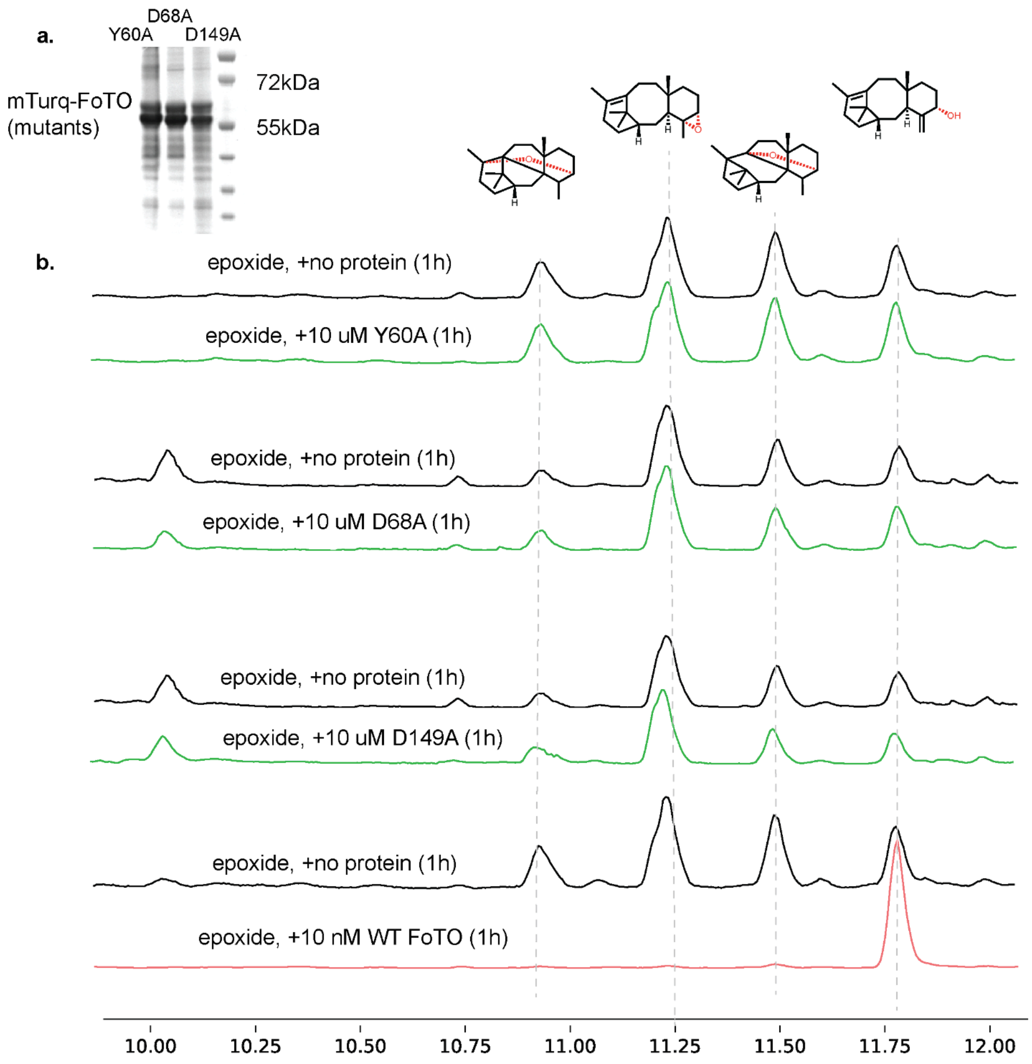


**Supplementary Fig 11**: **GC-MS analysis of the incubation of FoTO1 mutants with the epoxide intermediate**

**(a)** SDS-PAGE of elution fraction (eluted with 200mM imidazole) from bacterially-expressed FoTO1 (expected MW ~58kDa) & mutants after Ni-NTA column chromatography.

(**b**) GC-MS total ion chromatograms of purified FoTO1 (WT or point mutants as indicated) incubated with taxadiene-4(5)-epoxide in liposomes *in vitro*. TICs of three independent experiments.

**Table 1: All non-native genes infiltrated into *N. benthamiana* to be labelled by TurboID**

| Name | Notes | Amino Acid Sequence (AA) |
|---|---|---|
| **V5-TurboID** | Bait for control samples | MGKPIPNPLLGLDSTASKDNTVPLKLIALLANGEFHSGEQLGETLGMSRAAINKHIQT<br>LRDWGVDVFTVPGKGYSLPEPIPLLNAKQILGQLDGGSVAVLPVVDSTNQYLLDRIG<br>ELKSGDACIAEYQQAGRGSRGRKWFSPFGANLYLSMFWRLKRGPAAIGLGPVIGIV<br>MAEALRKLGADKVRVKWPNDLYLQDRKLAGILVELAGITGDAAQIVIGAGINVAMRRV<br>EESVVNQGWITLQEAGINLDRNTLAATLIRELRAALELFEQEGLAPYLPRWEKLDNFI<br>NRPVKLIIGDKEIFGISRGIDKQGALLLEQDGVIKPWMGGEISLRSAEK |
| **V5-TurboID-FOTO** | Bait for experimental samples | MGKPIPNPLLGLDSTASKDNTVPLKLIALLANGEFHSGEQLGETLGMSRAAINKHIQT<br>LRDWGVDVFTVPGKGYSLPEPIPLLNAKQILGQLDGGSVAVLPVVDSTNQYLLDRIG<br>ELKSGDACIAEYQQAGRGSRGRKWFSPFGANLYLSMFWRLKRGPAAIGLGPVIGIV<br>MAEALRKLGADKVRVKWPNDLYLQDRKLAGILVELAGITGDAAQIVIGAGINVAMRRV<br>EESVVNQGWITLQEAGINLDRNTLAATLIRELRAALELFEQEGLAPYLPRWEKLDNFI<br>NRPVKLIIGDKEIFGISRGIDKQGALLLEQDGVIKPWMGGEISLRSAEKGSGAETMNE<br>KVGADEKLIEDSRNTGQEENDYSKRLGPLSRNIIPHLINIYTCIATPRDLEIYHPDATFE<br>DFFVRAFGIKEIKSIHYSFPKLVYDGKILEYSVEENETSPGCGELLFNIKQQYKVLYLG<br>KEVNITTLMVLQIENGKIIKHEDRINQHPVRGRHDISVPLVGRAREGIRRLTMLMLHVR<br>MGFGKDPTPP |

| Protein | Amino Acid Sequence (AA) | Student's t-test Difference | Significance (-log10 p-value) |
|---|---|---|---|
| **TDS1** | AAC49310.1 | -0.03486 | 0.045088 |
| **TDS2** | De La Peña (2021) | 0.274253 | 0.299509 |
| **HMGR** | De La Peña (2021) | 0.565369 | 0.407609 |
| **chlGGPPS** | AAS49033.1 | 0.419902 | 1.411194 |
| **T5aH** | AAQ56240.2 | 2.377316 | 1.401159 |
| **T9dA** | McClune (2025) | -0.17731 | 2.326367 |
| **T1bH** | McClune (2025) | 0.809191 | 0.827117 |
| **PCL** | McClune (2025) | 0.496939 | 1.220664 |
| **TAT** | AAF34254.1 | -0.3256 | 0.680816 |
| **T10bH** | AAK00946.1 | 0.996758 | 2.170581 |
| **T13aH** | AAL23619.1 | 2.120403 | 1.85029 |
| **DBAT** | AAL57617.1 | 0.431275 | 0.324826 |

| | | | |
|---|---|---|---|
| **T2aH** | AAS89065.2 | 1.259913 | 0.979673 |
| **T7bH** | AAQ75553.1 | 1.050815 | 1.051024 |
| **TOT** | MDRVREIFNGSSGSPAGIPHSVITAGVGA<br>IIIILLSLLLLRRSSKRGDSSHPPGNSGLPF<br>IGETLSFTKAFKSNTLAEFFEERVKKFGN<br>VFKISIIGPPTVVMCGNEGNRFIFANEEKL<br>VHLSWSGRYAKILGGDSVSMKRGDDHR<br>SVRAAFAGFLSPASLPIYISKMSAQIQDHI<br>NQKWKGKDVIAVVPLVKELVFNVSYNLFF<br>SINDSEELHRLHKIFETIVEGHVSMPIDLP<br>GFTFHRALQGRAKLKKVFSSLIERRRSDL<br>SSGLASANQDLISVLLTYKDDRGYTMTH<br>DELLDNFLSLLESSYDSVNSPMACIFKLL<br>YANPECYEKVVQEQLGILSGKKEGQEIS<br>WKDLRSMKYTWQVLQETLRLYTQVAGIF<br>RKAMTVIHYDGHTIPKGWQLLWATQTTH<br>LNDKYFSEPEKFMPSRFDEEGNNVIPYS<br>FVPFGGGRRMCPGWEFGKMEILLFVHH<br>FVKAFSGFTPIDPDEKITGNPFPHLPANG<br>FSIKPTPRS | 0.99669075 | 0.688965 |
| **eGFP** | MTSKGEELFTGVVPILVELDGDVNGHKF<br>SVSGEGEGDATYGKLTLKFICTTGKLPVP<br>WPTLVTTFSYGVQCFSRYPDHMKRHDF<br>FKSAMPEGYVQERTIFFKDDGNYKTRAE<br>VKFEGDTLVNRIELKGIDFKEDGNILGHKL<br>EYNYNSHNVYIMADKQKNGIKVNFKIRH<br>NIEDGSVQLADHYQQNTPIGDGPVLLPD<br>NHYLSTQSALSKDPNEKRDHMVLLEFVT<br>AAGITHGMDELYK | 0.137591 | 0.143587 |
| **PAM** | AAT47186.1 | -0.66586 | 1.687681 |
| **DBTNBT** | AAM75818.1 | -0.11904 | 0.302004 |
| ***Cs*CYP71CD1** | KDO78504.1 | 2.039624 | 2.12514 |
| ***At*KAO** | ABJ17103.1 | 0.480021 | 0.399775 |

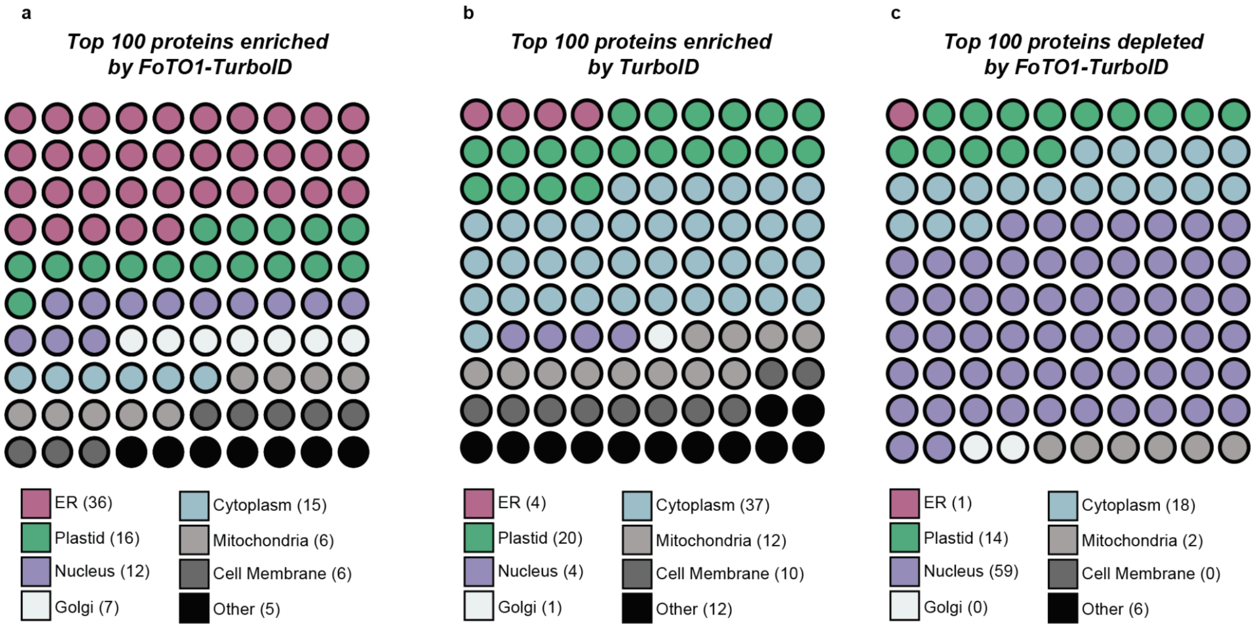


**Supplementary Figure 12: Localization of top TurboID control hits**

(a) Dot plot displaying the localization of the top 100 most enriched endogenous *N. benthamiana* proteins in the FoTO1-TurboID screen by FoTO1-TurboID

(b) Dot plot displaying the localization of the top 100 most enriched endogenous *N. benthamiana* proteins in the FoTO1-TurboID screen by TurboID.

(c) Dot plot displaying the localization of the top 100 most depleted endogenous *N. benthamiana* proteins in the FoTO1-TurboID screen by comparing FoTO1-TurboID hits to TurboID hits.

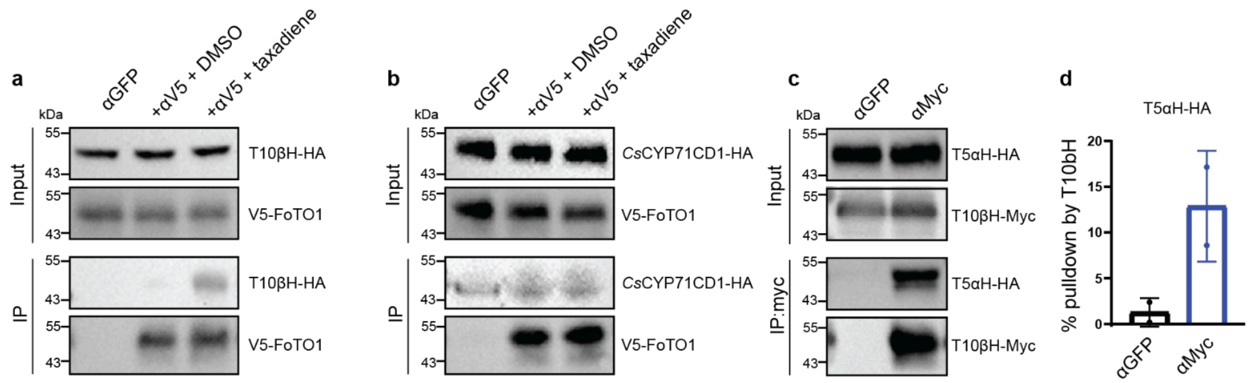


**Supplementary Figure 13: Co-immunoprecipitation of T5aH by T10bH in the absence of FoTO1**

(**a-b**) Representative immunoblot analysis of the co-IP of (**a**) T10βH-HA, (**b**) citrus CYP71CD1-HA by pulldown with V5-FoTO1 in co-infiltrated *N. benthamiana* leaves. αGFP lanes indicate internal negative control pulldowns where a GFP antibody was used for pulldown instead of a V5 antibody. Representative images of two independent experiments.

(**c**) Representative immunoblot analysis of the co-IP of T5αH-HA by pulldown with T10βH-Myc in co-infiltrated *N. benthamiana* leaves. αGFP lanes indicate internal negative control pulldowns where a GFP antibody was used for pulldown instead of a Myc antibody. Representative images of two independent experiments.

(**d**) Quantification of the immunoblot analysis of the co-immunoprecipitation (co-IP) of T5αH-HA by T10βH-myc. Values ± SD of two independent experiments ($*p \le 0.05$ by Student's t test and not shown if insignificant ($p > 0.05$)).

**a**

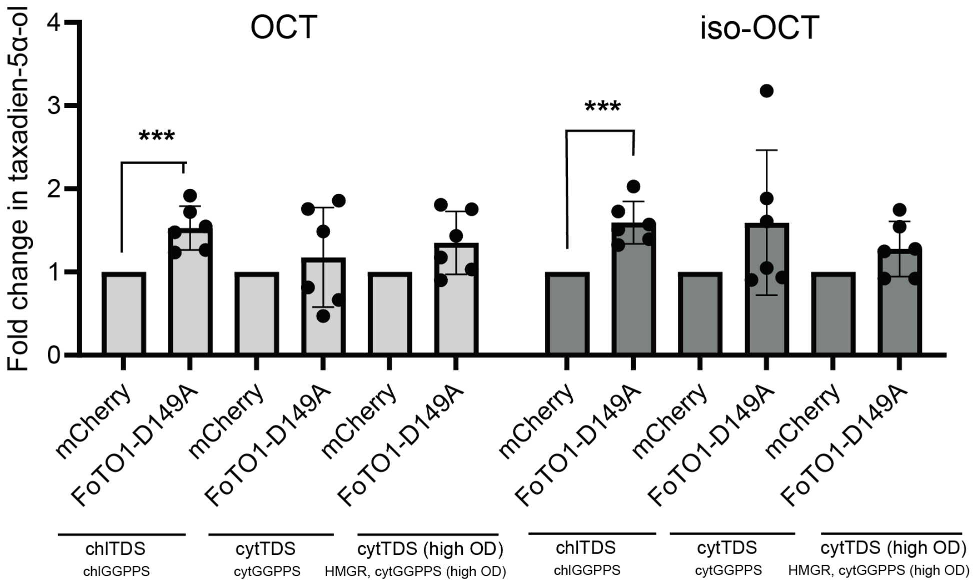


**Supplementary Fig 14: FoTO1 has an outsized effect when a large amount of TDS is infiltrated**

(**a**) Bar graph showing integrated peak area of OCT and iso-OCT by GC-MS when mCherry (ctrl) and FoTO1-D149A are transiently expressed in *N. benthamiana* leaves along with boost and early pathway enzymes (HMGR, GGPPS, TDS1, TDS2, T5αH) under the indicated conditions where standard OD is 0.2 and a high OD is 1.0. Ctrl, mCherry is used instead of FoTO1 as a negative control and wild-type FoTO1 is used as a positive control. One side of the leaf was infiltrated with a mix containing mCherry and the other side was infiltrated with a mix containing FoTO1-D149A. Fold change is calculated as the ratio of the measured peak area of taxadien-5α-ol on the FoTO1-D149A side of the leaf divided by the measured peak area of leaf tissue from the control side of the leaf. Values ± SD of six independent experiments (∗p ≤ 0.05, ∗*∗p ≤ 0.005 by Student's t test and not shown if insignificant (p > 0.05)).

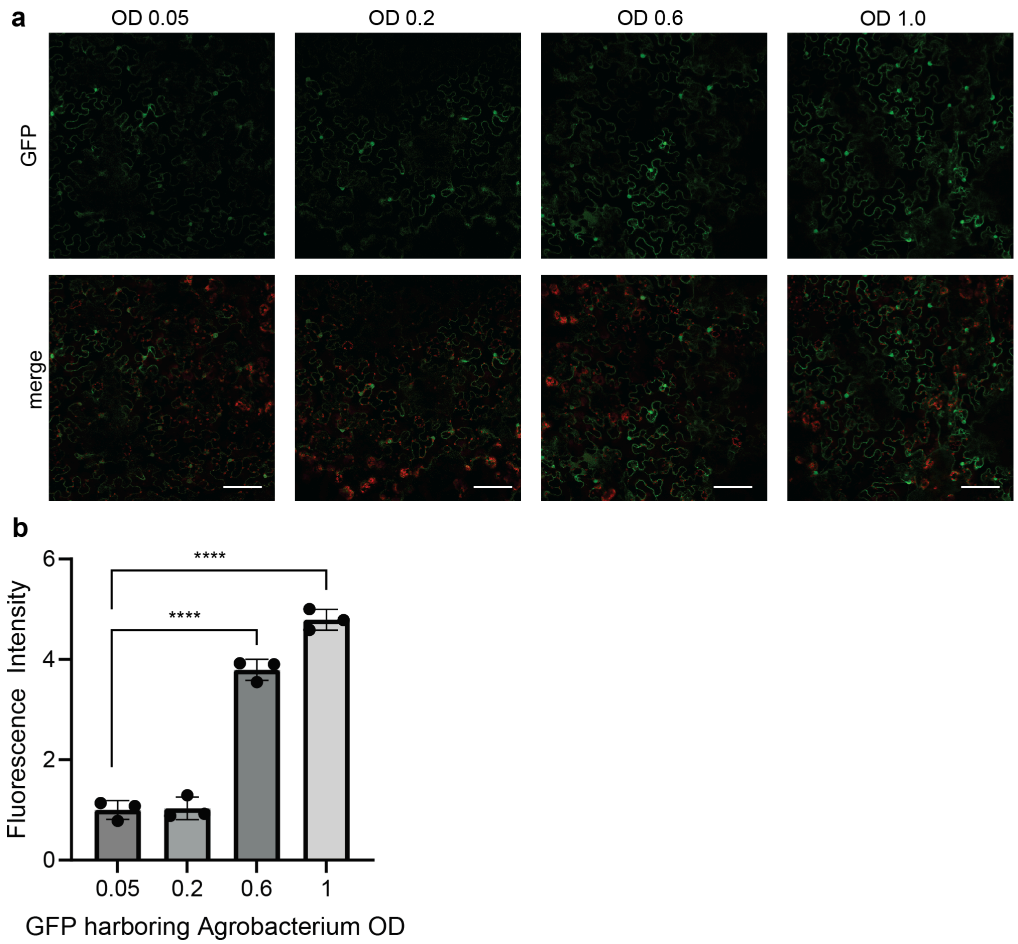


**Supplementary Fig 15: Increasing the OD of GFP harboring *Agrobacterium* increases the expression of GFP**

(**a**) Confocal microscopy images of *N. benthamiana* leaves infiltrated with *Agrobacterium* strains harboring GFP at the indicated ODs two days post-infiltration. GFP signal is pseudocolored in green and chloroplast autofluorescence is shown in red; merge is the overlay of the chloroplast autofluorescence (the scale bar represents 100 μm). Representative images of 3 independent experiments.

(**b**) Quantification of GFP fluorescence as observed by confocal microscopy when the indicated ODs of *Agrobacterium* harboring GFP are infiltrated into *N. benthamiana*. Values ± SD of three experiments (****$p \leq 0.0001$ by Student's t test and not shown if insignificant ($p > 0.05$)).

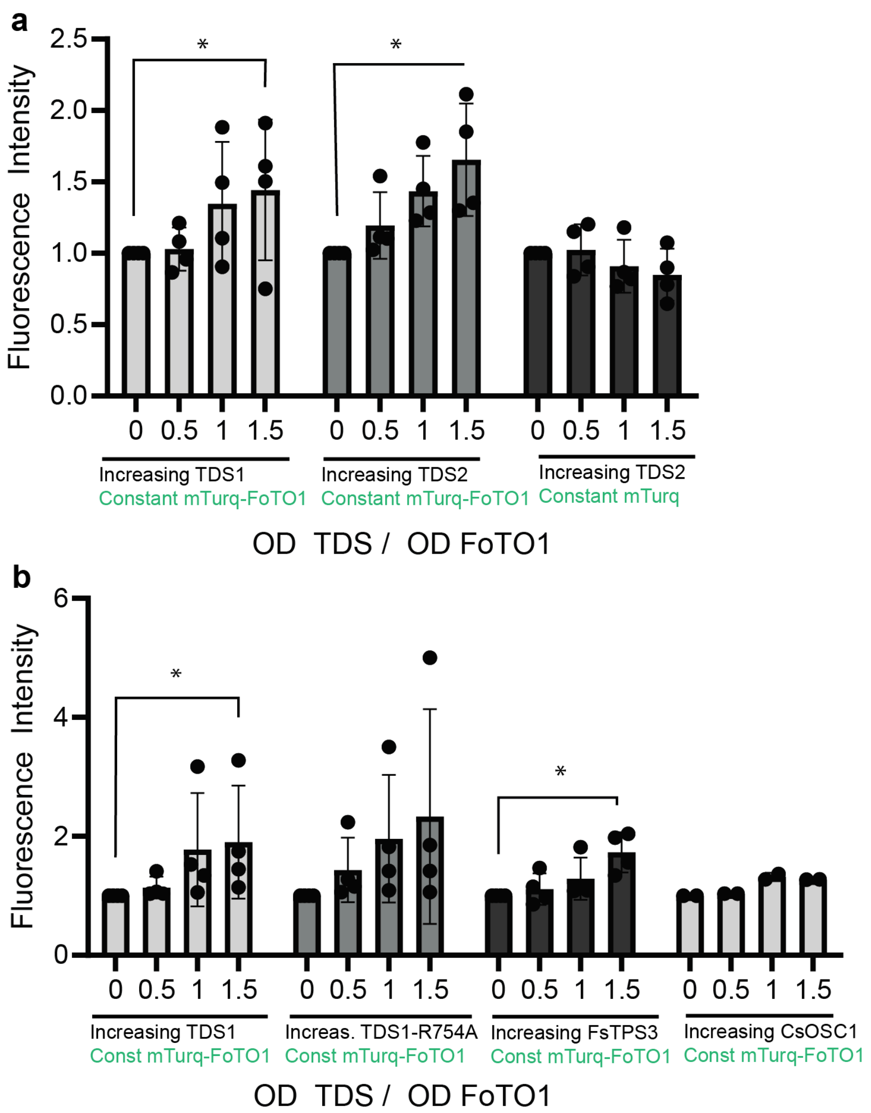


**Supplementary Figure 16: Increasing the amount of TDS increases the apparent expression of FoTO1**

(**a-b**) Bar graph showing mean fluorescence intensity of leaf areas co-infiltrated with increasing amounts of the indicated *Agrobacterium* strains and a constant OD of either mTurq-FoTO1 or mTurq strains as indicated. mCherry is used as a filler strain to balance the OD of the increasing strain. Values ± SD of four experiments (****$p \leq 0.0001$ by ordinary one-way ANOVA and not shown if insignificant ($p > 0.05$)).

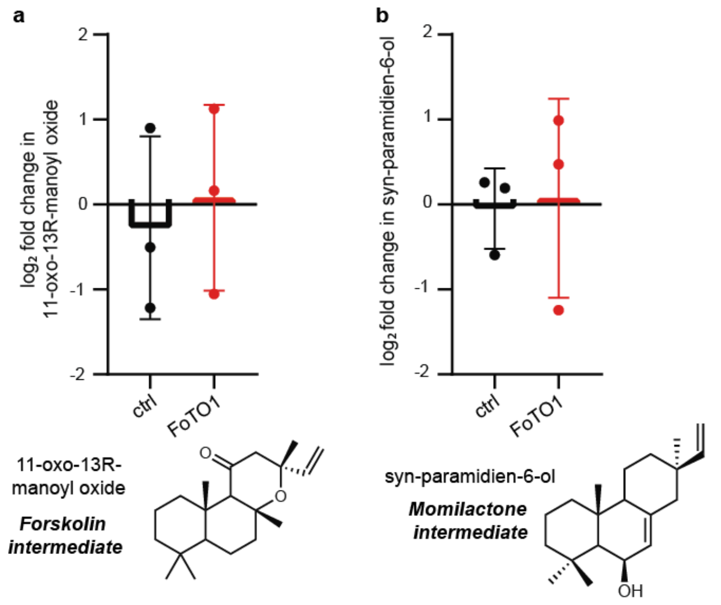


**Supplementary Figure 17: FoTO1 has a diminished effect on diverse cytoplasmic diterpene pathways**

(**a**) Quantification of the log2 fold change in 11-oxo-13R-manoyl oxide when cytoplasmic forskolin early pathway enzymes (cytGGPPS, cyt*Fs*TPS2, cyt*Fs*TPS3, *Fs*CYP71AH15) are transiently expressed in *N. benthamiana* leaves with and without FoTO1.

(**b**) Quantification of the log2 fold change in syn-paramidine-6-ol when cytoplasmic momilactone early pathway enzymes (cytGGPPS, cyt*Os*KSL4, cyt*Os*CPS4, *Os*CYP76M8) are transiently expressed in *N. benthamiana* leaves with and without FoTO1.

Values ± SD of three experiments (*$p \leq 0.05$ by Student's t-test and not shown if insignificant ($p > 0.05$)).

### Table 2: Mass Spectra of Compounds in this Study

| Compound | Molecular Formula | Calc. Mass | Retention Time | Mass Spectra |
|---|---|---|---|---|
| taxadiene (**1**) | $C_{20}H_{32}$ | 272.5 | 10.27 (GCMS) | 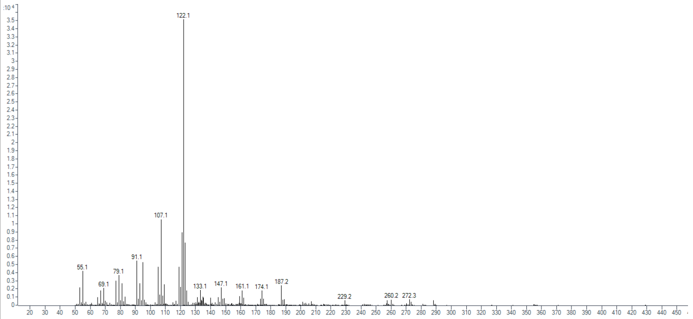 |
| taxadien-5a-ol (**2**) | $C_{20}H_{32}O$ | 288.5 | 11.81 (GCMS) | 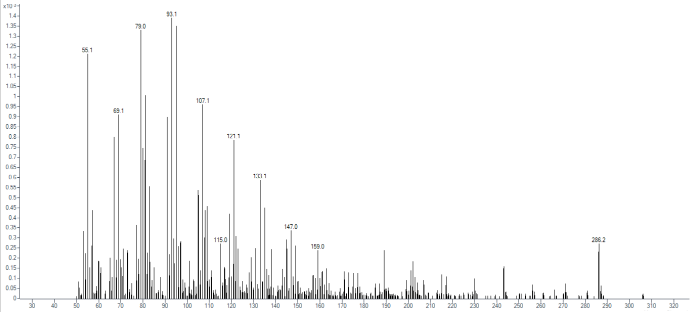 |
| OCT (**2a**) | $C_{20}H_{32}O$ | 288.5 | 11.53 (GCMS) | 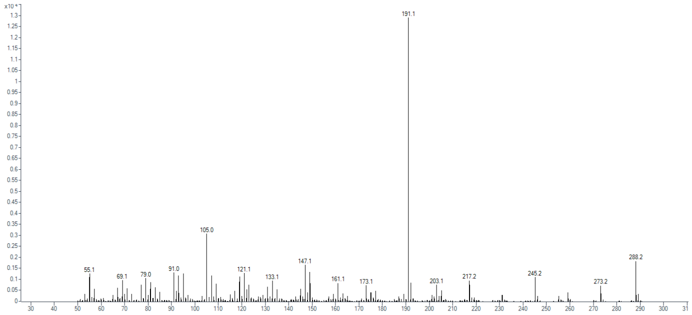 |
| iso-OCT (**2b**) | $C_{20}H_{32}O$ | 288.5 | 10.97 (GCMS) | 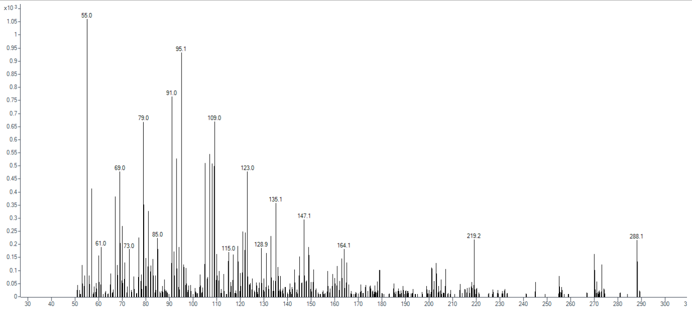 |

| | | | | |
|---|---|---|---|---|
| **2d** | $C_{20}H_{32}O$ | 288.5 | 12.67 (GCMS) | 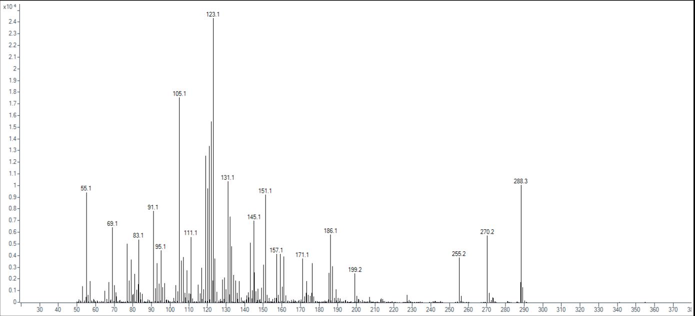 |
| 13R-manoyl oxide | $C_{20}H_{34}O$ | 290.5 | 10.02 (GCMS) | 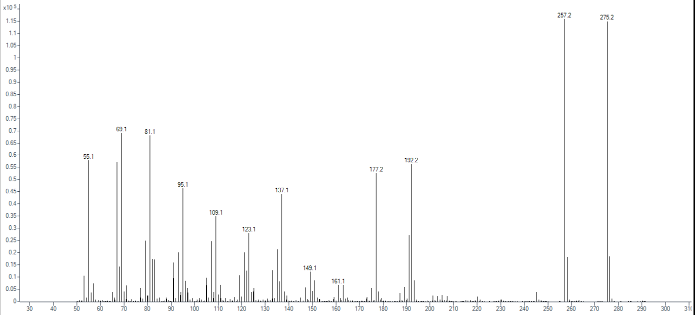 |
| 11-oxo-manoyl oxide | $C_{20}H_{32}O_2$ | 304.5 | 10.76 (GCMS) | 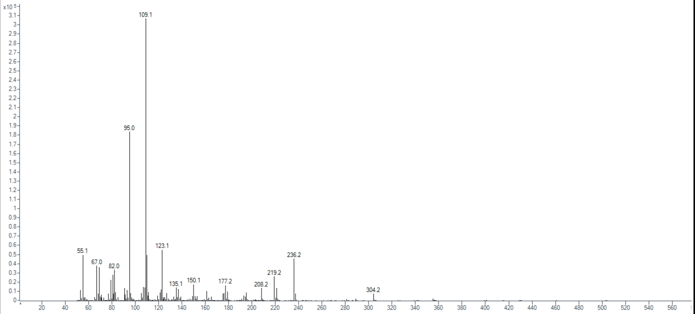 |
| syn-pimara diene | $C_{20}H_{32}$ | 272.5 | 12.34 (GCMS) | 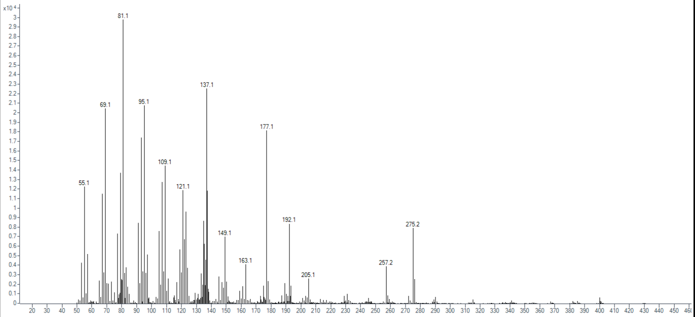 |
| syn-pimara dien-6-ol | $C_{20}H_{32}O$ | 288.5 | 11.58 (GCMS) | 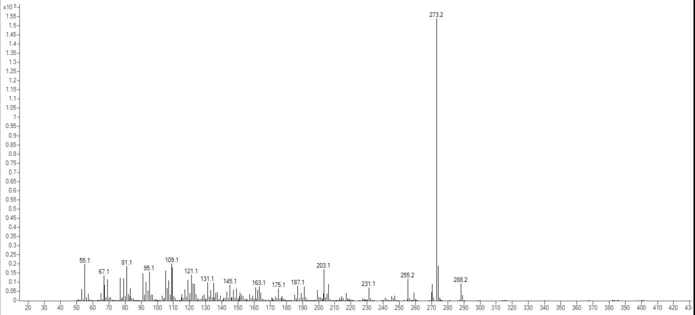 |

| | | | | |
|---|---|---|---|---|
| dihydro niloticin | $C_{30}H_{50}O_3$ | 458.7 | 12.12 (GCMS) | 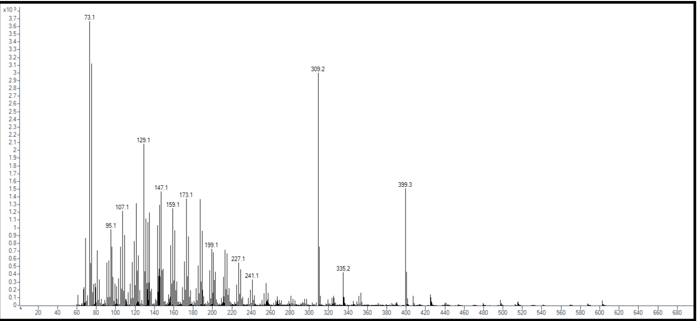 |
| $GA_{12}$ | $C_{20}H_{28}O_4$ | 332.4 | 8.76 (LCMS) | 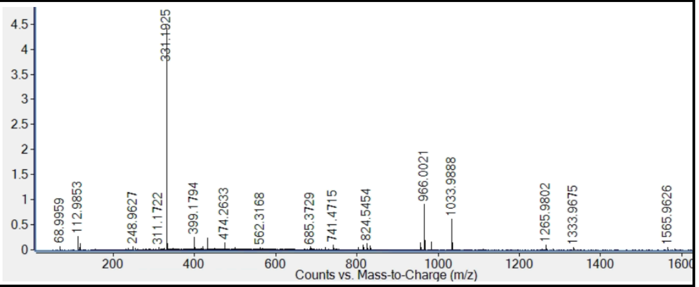 |

### Table 3: Primers Used in this Study

| Plasmid | Direction | Fragment | Sequence | Template |
|---|---|---|---|---|
| pCold-mTurq-FoTO1strain | Fwd | 1 | GCAGAAACAATGAATGAGAAAGTTGGC | pEAQ-FoTO1 |
| | Rev | 1 | CTATGGAGGAGTTGGATCCTTTCCAA | |
| pYeDP60-mTurq-FoTO1 FoTO1 yeast expression plasmid (URA3, ADE2)<br><br>pYeDP60 linearized with SacII and EcoRI | Fwd | 1 | tacacacactaaattaccggatccgaattcATGGTGAGTAAGGGTGAAGAATTATTTAC | pEAQ-mTurq-FoTO1 (previous work) |
| | Rev | 1 | tagagacatgggagatcccccgcggaattcCTATGGAGGAGTTGGATCCTTTCCAAAGCC | |
| FoTO1 expression plasmid for yeast expression<br><br>(The four fragments are ligated together) | Fwd | 1 | gaggcaagctaaacagatctgatcaaaaatcatcgcttcgc | pYeDP60-mTurq-FoTO1 |
| | Rev | 1 | ggtgacactatagaacgcggaacgcagaattttcgagtt | |
| | Fwd | 2 | cgagtcagtgagcgaggaataatggtttcttagtatgatccaatatcaaag | pYeDP60 |
| | Rev | 2 | ctgccggtctccgaaaagtgccacctgaacgaagcatctgtg | |
| | Fwd | 3 | aactcgaaaattctgcgttccgcgttctatagtgtcacc | pFA6a-kanMX6 |
| | Rev | 3 | ctttgatattggatcatactaagaaaccattattcctcgctcactgactcg | |
| | Fwd | 4 | cacagatgcttcgttcaggtggcacttttcggagaccggcag | pFA6a-kanMX6 |
| | Rev | 4 | gcgaagcgatgatttttgatcagatctgtttagcttgcctc | |
| pEAQ-V5-FoTO1-D68A | Fwd | 1 | AGCTTCTGTATATTCTGCCCAAATTCGCGACCGGT | pEAQ-V5-FoTO1 (previous work) |
| | Rev | 1 | CACGCACAAAGAAGGCTTCAAATGTCGCAT | |
| | Fwd | 2 | ATGCGACATTTGAAGCCTTCTTTGTGCGTG | pEAQ-V5-FoTO1 |
| | Rev | 2 | GAAAATTTAATGAAACCAGAGTTAAAGGCCTCGAGCTATGGAGGAGTTGGATCCTTTCCA | |
| pEAQ-V5-FoTO1-D149A | Fwd | 1 | AGCTTCTGTATATTCTGCCCAAATTCGCGACCGGT | pEAQ-V5-FoTO1 |
| | Rev | 1 | TGTTGGTTAATCCGGGCTTCGTGCTTGATG | |
| | Fwd | 2 | CATCAAGCACGAAGCCCGGATTAACCAACA | pEAQ-V5-FoTO1 |
| | Rev | 2 | GAAAATTTAATGAAACCAGAGTTAAAGGCCTCGAGCTATGGAGGAGTTGGATCCTTTCCA | |
| pEAQ-V5-FoTO1-Y60A | Fwd | 1 | AGCTTCTGTATATTCTGCCCAAATTCGCGACCGGT | pEAQ-V5-FoTO1 |
| | Rev | 1 | TCGCATCCGGATGAGCTATTTCAAGGTCACG | |

| | | | | |
|---|---|---|---|---|
| | Fwd | 2 | CGTGACCTTGAAATAGCTCATCCGGATGCGA | pEAQ-V5-FoTO1 |
| | Rev | 2 | GAAAATTTAATGAAACCAGAGTTAAAGGCCTCGAGCTATGGAGGAGTTGGATCCTTTCCA | |
| pEAQ-V5-FoTO1-Y60S | Fwd | 1 | AGCTTCTGTATATTCTGCCCAAATTCGCGACCGGT | pEAQ-V5-FoTO1 |
| | Rev | 1 | ATCCGGATGAGATATTTCAAGGTCA | |
| | Fwd | 2 | CCTTGAAATATCTCATCCGGATGC | pEAQ-V5-FoTO1 |
| | Rev | 2 | GAAAATTTAATGAAACCAGAGTTAAAGGCCTCGAGCTATGGAGGAGTTGGATCCTTTCCA | |
| pEAQ-V5-FoTO1-Y60F | Fwd | 1 | AGCTTCTGTATATTCTGCCCAAATTCGCGACCGGT | pEAQ-V5-FoTO1 |
| | Rev | 1 | ATCCGGATGAAATATTTCAAGGTCA | |
| | Fwd | 2 | CCTTGAAATATTTCATCCGGATGC | pEAQ-V5-FoTO1 |
| | Rev | 2 | GAAAATTTAATGAAACCAGAGTTAAAGGCCTCGAGCTATGGAGGAGTTGGATCCTTTCCA | |
| pEAQ-V5-FoTO1-Y60H | Fwd | 1 | AGCTTCTGTATATTCTGCCCAAATTCGCGACCGGT | pEAQ-V5-FoTO1 |
| | Rev | 1 | ATCCGGATGATGTATTTCAAGGTCA | |
| | Fwd | 2 | CCTTGAAATACATCATCCGGATGC | pEAQ-V5-FoTO1 |
| | Rev | 2 | GAAAATTTAATGAAACCAGAGTTAAAGGCCTCGAGCTATGGAGGAGTTGGATCCTTTCCA | |
| pEAQ-V5-FoTO1-D68E | Fwd | 1 | AGCTTCTGTATATTCTGCCCAAATTCGCGACCGGT | pEAQ-V5-FoTO1 |
| | Rev | 1 | CGCACAAAGAATTCTTCAAATGTC | |
| | Fwd | 2 | GACATTTGAAGAATTCTTTGTGCG | pEAQ-V5-FoTO1 |
| | Rev | 2 | GAAAATTTAATGAAACCAGAGTTAAAGGCCTCGAGCTATGGAGGAGTTGGATCCTTTCCA | |
| pEAQ-V5-FoTO1-D68N | Fwd | 1 | AGCTTCTGTATATTCTGCCCAAATTCGCGACCGGT | pEAQ-V5-FoTO1 |
| | Rev | 1 | CGCACAAAGAAGTTTTCAAATGTC | |
| | Fwd | 2 | GACATTTGAAAACTTCTTTGTGCG | pEAQ-V5-FoTO1 |
| | Rev | 2 | GAAAATTTAATGAAACCAGAGTTAAAGGCCTCGAGCTATGGAGGAGTTGGATCCTTTCCA | |
| pEAQ-V5-FoTO1-D68H | Fwd | 1 | AGCTTCTGTATATTCTGCCCAAATTCGCGACCGGT | pEAQ-V5-FoTO1 |
| | Rev | 1 | CGCACAAAGAAGTGTTCAAATGTC | |

| | | | | |
|---|---|---|---|---|
| | Fwd | 2 | GACATTTGAACACTTCTTTGTGCG | pEAQ-V5-FoTO1 |
| | Rev | 2 | GAAAATTTAATGAAACCAGAGTTAAAGGCCTCGAGCTATGGAGGAGTTGGATCCTTTCCA | |
| pEAQ-V5-FoTO1-D149E | Fwd | 1 | AGCTTCTGTATATTCTGCCCAAATTCGCGACCGGT | pEAQ-V5-FoTO1 |
| | Rev | 1 | GTTAATCCGTTCTTCGTGCTT | |
| | Fwd | 2 | AAGCACGAAGAACGGATTAAC | pEAQ-V5-FoTO1 |
| | Rev | 2 | GAAAATTTAATGAAACCAGAGTTAAAGGCCTCGAGCTATGGAGGAGTTGGATCCTTTCCA | |
| pEAQ-V5-FoTO1-D149N | Fwd | 1 | AGCTTCTGTATATTCTGCCCAAATTCGCGACCGGT | pEAQ-V5-FoTO1 |
| | Rev | 1 | GTTAATCCGGTTTTCGTGCTT | |
| | Fwd | 2 | AAGCACGAAAACCGGATTAAC | pEAQ-V5-FoTO1 |
| | Rev | 2 | GAAAATTTAATGAAACCAGAGTTAAAGGCCTCGAGCTATGGAGGAGTTGGATCCTTTCCA | |
| pEAQ-V5-FoTO1-D149H | Fwd | 1 | AGCTTCTGTATATTCTGCCCAAATTCGCGACCGGT | pEAQ-V5-FoTO1 |
| | Rev | 1 | GTTAATCCGGTGTTCGTGCTT | |
| | Fwd | 2 | AAGCACGAACACCGGATTAAC | pEAQ-V5-FoTO1 |
| | Rev | 2 | GAAAATTTAATGAAACCAGAGTTAAAGGCCTCGAGCTATGGAGGAGTTGGATCCTTTCCA | |
| pEAQ-V5-TurboID-FoTO1 | Fwd | 1 | CGGTCTGCCGAAAAGGGCTCTGGAGCAGAAACA | pEAQ-V5-FoTO1 |
| | Rev | 1 | GAAAATTTAATGAAACCAGAGTTAAAGGCCTCGAGCTATGGAGGAGTTGGATCCTTTCCA | |
| | Fwd | 2 | GCCCAAATTCGCGACCGGCAAATTCGCGACCGGATGGGCAAGCCCATC | pEAQ-V5-TurboID (previous work) |
| | Rev | 2 | TGTTTCTGCTCCAGAGCCCTTTTCGGCAGACCG | |
| pEAQ-T10bH-HA | Fwd | 1 | AGCTTCTGTATATTCTGCCCAAATTCGCGACCGGT | pEAQ-T10bH (previous work) |
| | Rev | 2 | TTAAGCATAATCTGGAACATCATATGGATATCCAGAGCCGGATCTCGGAA | |
| | Rev | 2 | GAAACCAGAGTTAAAGGCCTCGAGTTAAGCATAATCTGGAACATCATATGGATATCCAGAGCC | |
| pEAQ-T10bH-myc | Fwd | 1 | AGCTTCTGTATATTCTGCCCAAATTCGCGACCGGT | pEAQ-T10bH |

| | | | | |
|---|---|---|---|---|
| | Rev | 2 | TCACAGATCCTCTTCTGAGATGAGTTTTTGTTCTCCAGAGCCGGATCTCGGAAAAAGTTTTAT | |
| | Rev | 2 | GAAACCAGAGTTAAAGGCCTCGAGTCACAGATCCTCTTCTGAGATGAGTTTTTGTTCTCCAGAGCC | |
| pEAQ-T13aH-HA | Fwd | 1 | AGCTTCTGTATATTCTGCCCAAATTCGCGACCGGT | pEAQ-T13aH (previous work) |
| | Rev | 2 | TTAAGCATAATCTGGAACATCATATGGATATCCAGAGCCAGATCTGGAATAGAGTTT | |
| | Rev | 2 | GAAACCAGAGTTAAAGGCCTCGAGTTAAGCATAATCTGGAACATCATATGGATATCCAGAGCC | |
| pEAQ-T2aH-HA | Fwd | 1 | AGCTTCTGTATATTCTGCCCAAATTCGCGACCGGT | pEAQ-T2aH (previous work) |
| | Rev | 2 | ATAATCTGGAACATCATATGGATATCCAGAGCCGGATCGAGAAATAAGTTT | |
| | Rev | 2 | GAAACCAGAGTTAAAGGCCTCGAGTTAAGCATAATCTGGAACATCATATGGATATCCAGAGCC | |
| pEAQ-*Cs*CYP71CD1-HA | Fwd | 1 | TATTCTGCCCAAATTCGCGACCGGTATGGAGCAACAATTTGATTAC | pEAQ-*Cs*CYP71CD1 (previous work) |
| | Rev | 2 | ATAATCTGGAACATCATATGGATATCCAGAGCCCGGAATATTATTGTAAGGAG | |
| | Rev | 2 | GAAACCAGAGTTAAAGGCCTCGAGTTAAGCATAATCTGGAACATCATATGGATATCCAGAGCC | |
| pEAQ-YFPn | Fwd | 1 | CTGCCCAAATTCGCGACCGGTATGGTGAGCAAGGGC | eYFP |
| | Rev | 1 | ACCAGAGTTAAAGGCCTCGAGCTAGTCCTCGATGTTGTG | |
| pEAQ-FoTO1-YFPn | Fwd | 1 | AGATCTATTGCTACTATGGTGAGCAAGGGC | eYFP |
| | Rev | 1 | ACCAGAGTTAAAGGCCTCGAGCTAGTCCTCGATGTTGTG | |
| | Fwd | 2 | AGCTTCTGTATATTCTGCCCAAATTCGCGACCGGT | pEAQ-FoTO1 |
| | Rev | 2 | AGTAGCAATAGATCTTGGAGGAGTTGGATC | |
| pEAQ-YFPc-FoTO1 | Fwd | 1 | CTGCCCAAATTCGCGACCGGTATGAAGCAGAAGAACGGC | eYFP |
| | Rev | 1 | AGTAGCAATAGATCTCTTGTACAGCTCGTC | |
| | Fwd | 2 | AGATCTATTGCTACTGCAGAAACAATGAAT | pEAQ-FoTO1 |
| | Rev | 2 | GAAAATTTAATGAAACCAGAGTTAAAGGCCTC | |

| | | | | |
|---|---|---|---|---|
| | | | GAGCTATGGAGGAGTTGGATCCTTTCCA | |
| pEAQ-Calnexin-YFPc | Fwd | 1 | CGCTCGATCGCGACGAAGCAGAAGAACGGCATCAAGG | eYFP |
| | Rev | 1 | GAAAATTTAATGAAACCAGAGTTAAAGGCCTCGAGTTACTTGTACAGCTCG | |
| | Fwd | 2 | CTTCTGTATATTCTGCCCAAATTCGCGACCGGATGGAGGAGCGGAGCCGG | *Nb* cDNA |
| | Rev | 2 | CGTCGCGATCGAGCGATTATCACGTCTCGT | |
| pEAQ-CPR-YFPc | Fwd | 1 | AGATCTATTGCTACTAAGCAGAAGAACGGC | eYFP |
| | Rev | 1 | GAAAATTTAATGAAACCAGAGTTAAAGGCCTCGAGTTACTTGTACAGCTCG | |
| | Fwd | 2 | AGCTTCTGTATATTCTGCCCAAATTCGCGACCGGT | pEAQ-CPR (previous work) |
| | Rev | 2 | AGTAGCAATAGATCTCCATATATCTCGTAAGT | |
| pEAQ-T5aH-YFPc | Fwd | 1 | AGATCTATTGCTACTAAGCAGAAGAACGGC | eYFP |
| | Rev | 1 | GAAAATTTAATGAAACCAGAGTTAAAGGCCTCGAGTTACTTGTACAGCTCG | |
| | Fwd | 2 | AGCTTCTGTATATTCTGCCCAAATTCGCGACCGGT | pEAQ-T5aH |
| | Rev | 2 | AGTAGCAATAGATCTTGGTCTCGGAAACAG | |
| pEAQ-TDS2-His | Fwd | 1 | AGCTTCTGTATATTCTGCCCAAATTCGCGACCGGT | pEAQ-TDS2 (previous work) |
| | Rev | 1 | CTAGTGGTGATGGTGATGATGTCCAGAGCCTACTTGAATCGGTTCAATGTAAACTTT | |
| | Rev | 1 | TTTAATGAAACCAGAGTTAAAGGCCTCGAGCTAGTGGTGATGGTGATGATGTCCAGAGCC | |
